# Ethylene signaling regulates natural variation in the abundance of antifungal acetylated diferuloylsucroses and *Fusarium graminearum* resistance in maize seedling roots

**DOI:** 10.1101/332056

**Authors:** Shaoqun Zhou, Ying K. Zhang, Karl A. Kremling, Yezhang Ding, John S. Bennett, Justin S. Bae, Dean K. Kim, Michael V. Kolomiets, Eric A. Schmelz, Frank C. Schroeder, Edward S. Buckler, Georg Jander

**Affiliations:** Boyce Thompson Institute, 533 Tower Road, Ithaca, NY 14853; Plant Biology Section, School of Integrated Plant Science, Cornell University, Ithaca, NY14853; Department of Chemistry and Chemical Biology, Cornell University, Ithaca, NY14853; Department of Plant Breeding and Genetics, Cornell University, Ithaca, NY14853; Section of Cell and Developmental Biology, University of California at San Diego, La Jolla, CA 92093; Department of Plant Pathology and Microbiology, Texas A&M University, College Station, TX, 77840; United States Department of Agriculture-Agricultural Research Service, Robert W. Holley Center for Agriculture and Health, Ithaca, New York 14853

**Keywords:** Acetylated diferuloylsucrose, ethylene, *Fusarium graminearum*, metabolite QTL mapping, Zea mays (maize)

## Abstract

- The production and regulation of defensive specialized metabolites plays a central role in pathogen resistance in maize (*Zea mays*) and other plants. Therefore, identification of genes involved in plant specialized metabolism can contribute to improved disease resistance.
- We used comparative metabolomics to identify previously unknown antifungal metabolites in maize seedling roots, and investigated the genetic and physiological mechanisms underlying their natural variation using quantitative trait locus (QTL) mapping and comparative transcriptomics approaches.
- Two maize metabolites, smilaside A (3,6-diferuloyl-3′,6′-diacetylsucrose) and smiglaside C (3,6-diferuloyl-2′,3′,6′-triacetylsucrose), that may contribute to maize resistance against *Fusarium graminearum* and other fungal pathogens were identified. Elevated expression of an ethylene receptor gene, *ETHYLENE INSENSITIVE 2* (*ZmEIN2*), co-segregated with decreased smilaside A/smiglaside C ratio. Pharmacological and genetic manipulation of ethylene availability and sensitivity *in vivo* indicated that, whereas ethylene was required for the production of both metabolites, the smilaside A/smiglaside C ratio was negatively regulated by ethylene sensitivity. This ratio, rather than the absolute abundance of these two metabolites, was important for maize seedling root defense against *F. graminearum.*
- Ethylene signaling regulates the relative abundance of the two *F. graminearum*-resistance-related metabolites and affects resistance against *F. graminearum* in maize seedling roots.

## Introduction

Plants in natural and manmade ecosystems are continuously exposed to microbial pathogens. Specialized metabolic pathways that give rise to diverse arsenals of bioactive defense compounds allow plants to efficiently fend off pathogen attacks. The significance of plant specialized metabolism in agriculture is exemplified by the association of specific biosynthetic genes with resistance against insect pests and phytopathogens (Meihls *et al.*, 2013; Handrick *et al.*, 2016; Yang *et al.*, 2017). Such studies highlight the potential of enlisting naturally occurring specialized metabolites in crop species to enhance quantitative disease resistance.

In North America, maize (*Zea mays*) is the most important agricultural crop, with over 13 billion bushels produced per annum, of which approximately 10% is lost to disease (Mueller, 2016a; Mueller, 2016b; Mueller, 2017). Maize is also known for its great genetic diversity, involving both nucleotide polymorphisms and structural genomic variation (Buckler *et al.*, 2006; Jiao *et al.*, 2017). The genetic architecture of disease resistance in maize has been investigated extensively using publicly available genetic resources (Mideros *et al.*, 2012; Olukolu *et al.*, 2014; Benson *et al.*, 2015). Compared to foliar and ear diseases, maize seedling diseases remain a relatively understudied area, even though in some years they can account for more yield loss than any single disease in the aboveground tissues (Mueller, 2016a). This may be due to the fact that experimental methods developed for large scale screening of diseases in aboveground tissues, such as controlled pathogen inoculation and visual symptom scoring, are difficult to apply to seedling diseases under field conditions.

*Fusarium graminearum* is one of the most common causal pathogens of maize seedling disease in the northern temperate zone. In the field, it overwinters on crop residue as thickened hyphae and produces asexual conidia that infect germinating seedlings roots or mesocotyls. Depending on the developmental stage and infection site, *F. graminearum* can also cause root rot, stem rot, and ear rot in maize (Munkvold & White, 2016). Previous research on maize-*F. graminearum* interactions has primarily focused on ear rot, with results consistently suggesting that resistance against this disease is most likely controlled by numerous small-effect quantitative trait loci (QTL) that are influenced by experimental methods and genetic backgrounds (Ali *et al.*, 2005; Kebede *et al.*, 2016; Brauner *et al.*, 2017). Furthermore, transcriptomic studies in maize and wheat show that host-*F. graminearum* interactions are significantly influenced by host tissue types, suggesting that the QTL associated with *F. graminearum* ear rot resistance probably will not confer resistance in seedling roots (Kazan *et al.*, 2012; Zhang *et al.*, 2016).

Compared to QTL identified from *F. graminearum* ear rot studies, factors contributing to *F. graminearum* stalk rot resistance may be more relevant to infections of seedling roots. For instance, near-isogenic lines (NILs) selected based on stalk rot resistance phenotypes also showed significant differences in primary root symptoms after controlled inoculation (Ye *et al.*, 2013). *Fusarium graminearum* infection in maize stalks induces production of specialized metabolites with antifungal activities (Huffaker *et al.*, 2011; Schmelz *et al.*, 2011). Additionally, comparative and correlative studies have identified constitutive phytoanticipins that were associated with *F. graminearum* resistance. For example, an *F. graminearum-*resistant NIL was found to accumulate significantly higher phenolic acids in its seedling roots compared to susceptible relatives. Interestingly, these differences disappear after *F. graminearum* infection, primarily due to fungus-induced reduction of defenses in the resistant NILs (Ye *et al.*, 2013). The same compounds also have been identified as metabolites related to *F. graminearum* resistance in other crop species, and were shown to inhibit fungal growth *in vitro* (Bollina *et al.*, 2010; Ponts *et al.*, 2011). Taken together, these studies indicate that specialized metabolites in maize seedling roots play a significant role in resistance against *F. graminearum.*

Here we describe a comparative metabolomics approach using maize inbred lines B73 and Mo17 to identify previously unknown maize antifungal compounds. These experiments led to the identification of two acetylated diferuloylsucroses, one of which demonstrated significant fungal growth inhibition *in vitro* at a physiologically relevant concentration. Genetic mapping, analysis of mutants, and physiological experiments demonstrated that accumulation of acetylated diferuloylsucroses is promoted by ethylene production and fine-tuned by ethylene sensitivity in maize.

## Materials and Methods

### Plant growth and fungal inoculation

All maize lines were obtained from the Maize Genetics Cooperation Stock Center (Urbana Champaign, Illinois). Maize seeds were germinated in moisturized rolls of germination paper, and seedlings were transplanted to 7.5 cm × 7.5 cm plastic pots with Turface^®^ MVP^®^ calcined clay (Profile Products LLC, Buffalo Grove, IL) when their primary roots reached approximately 9 cm. For inoculation experiments, seedling roots were immersed in *F. graminearum* spore suspension or mock solution (0.03% Phytagar) for one hour prior to transplanting. Spore suspensions were freshly prepared by flooding and scraping 7-day-old fungal cultures, maintained on potato dextrose agar plates, with 0.03% Phytagar suspension. The spore concentration was adjusted to be 5 × 10^5^ spores per ml after measurement with a hemocytometer and light microscopy. Hyphal fragments also were observed in the spore suspension. In all seedling inoculation experiments, *F. graminearum* strain ZTE, which was obtained from Dr. Frances Trail (Guenther & Trail, 2005), which was derived from a field-collected strain Z-3639 (Proctor *et al.*, 1995) transformed with the plasmid pTEFEGFP (Vanden Wymelenberg *et al.*, 1997), with permission from Dr. Robert Proctor. Transplanted seedlings were maintained in long day (16 hours) growth condition at 26 °C and 70% relative humidity. Six days after inoculation, the total root length of seedlings was measured with the RootReader 2D system (Famoso *et al.*, 2010). For screening of natural variation in *F. graminearum*-induced morphological changes, at least five seedlings of each plant line were mock- or fungus-inoculated in two independent experiments for root length measurement. In two other experiments, seven B73 and Mo17 seedlings were inoculated as described above, root length was measured, and tissue was harvested for further analysis.

### *Fusarium graminearum* gene expression measurement, mycotoxin quantification, and visual symptom scoring

The expression of the *F. graminearum* β-tubulin gene in seedling roots was measured using a previously published protocol (Lou *et al.*, 2016). The primers used for qRT-PCR detection of *F. graminearum* are FgTub-qF293 (5’-ATGCTTCCAACAACTATGCT-3’) and FgTub-qR411 (5’-AACTAGGAAACCCTGGAGAC-3’), which were designed based on the *F. graminearum* strain PH-1 reference genome sequence (Cuomo *et al.*, 2007). Fungal gene expression in each technical replicate was normalized by measurement of the expression of a constitutive maize actin gene with the following primers ZmActin-qF (5’-CCATGAGGCCACGTACAACT-3’) and ZmActin-qR (5’-GGTAAAACCCCACTGAGGA-3’).

Approximately 100 mg of fresh frozen seedling root tissue of each sample was used for extraction of deoxynivalenol, the main *F. graminearum* mycotoxin. Three microliters of 50:49.9:0.1 methanol:water:formic acid extraction solvent (Sigma-Aldrich) were added per mg of tissue. Ground tissue and extraction solvent were mixed and incubated at 4 °C for 40 min. Solid debris was separated from the solvent by centrifuging at 10,000 x *g* for 10 min. For each sample, 200 µL clear extract were filtered through a 0.45 micron filter plate by. Deoxynivalenol content in the extract was then measured with an Enzyme-Linked Immuno-Sorbent Assay (ELISA) kit following the manufacturer’s protocol (Helica Biosystems, Santa Ana, CA). Visual symptoms in seedling roots were scored for in three other comparative experiments with B73 and Mo17 seedlings 3 weeks after *F. graminearum* inoculation. In both experiments, at least 10 seedling roots of either genotype were scored on a 0-4 scale (0 = symptom-less; 1 = single restricted necrosis spot; 2 = single extended necrosis or multiple restricted necrosis spots; 3 = widespread necrosis throughout; 4 = seedling dead).

### Maize root metabolomics analyses

The extraction protocol for deoxynivalenol also was used for plant specialized metabolite extraction. Root extracts were analyzed using liquid chromatography-mass spectrometry (LC-MS). Chromatography was performed on a Dionex 3000 Ultimate UPLC-diode array detector system coupled to Thermo Q-Exactive mass spectrometer. Root extract samples were separated on a Titan C18 7.5 cm ×2.1 mm 1.9 μm, Supelco Analytical Column (Sigma-Aldrich) with a flow rate of 0.5 ml/min, using a gradient flow of 0.1 % formic acid in LC-MS grade water (eluent A) and 0.1 formic acid in acetonitrile (eluent B). Initial metabolite profiling experiments involved an 8-minute linear gradient from 95:5 A:B to 0:100 A:B for comparison of *F. graminearum* induced metabolomic changes in B73 and Mo17 seedling roots. This method was extended to a 15-minute gradient for better separation in later experiments. Mass spectral parameters were set as follows: Spray voltage 3500 V, Capillary temperature 300°C, Sheath Gas 35 units, Auxiliary Gas 10 units, probe heater temp 200°C with a HESI probe. Full scan mass spectra were collected (R:35000 FWHM at m/z 200; mass range: m/z 100 to 900) in both positive and negative ion spray modes. For non-targeted metabolomics analyses, metabolite abundance was estimated with signal intensity acquired through the XCMS-CAMERA mass scan data processing pipeline (Tautenhahn *et al.*, 2008; Benton *et al.*, 2010; Kuhl *et al.*, 2012). Smilaside A and smiglaside C abundance was estimated using peak areas at the respective *m/z* channel under negative electron spray ionization mode. Metabolite quantification was normalized by the total ion concentration to account for technical variation between samples. For non-targeted comparative metabolomics analyses, Student’s *t*-tests were used to identify mass features that were constitutively different between B73 and Mo17 (p<0.01, and fold change >2 or 0.01<p<0.05 and fold change >4) and significantly changed by *F. graminearum* in B73 (p<0.01, and fold change >1.5 or 0.01<p<0.05 and fold change >4).

### Structural identification of smiglaside C and smilaside A

To determine the chemical structures of smiglaside C and smilaside A, the LC-MS method for non-targeted metabolomics analysis was adapted to extract bulk maize seedling roots with 100% methanol at 4 °C overnight. Solid debris was filtered out and the crude extract was concentrated with a Buchi Rotovapor. The concentrated crude extract was fractionated on a normal phase column with a methanol:dichloromethane gradient on a CombiFlash Rf+ (Teledyne Isco, Lincoln, NE), and further purified for the target compounds with a water:acetonitrile gradient on a ZORBAX Eclipse XDB C18 column on an Agilent 1100 HPLC system (Agilent, Santa Clara, CA). Purified compounds were dried, weighed, and re-dissolved in methanol. NMR spectroscopy analyses were carried out on a Unity INOVA 600 instrument (Varian Medical Systems, Palo Alto, CA) with the following conditions: 256 scan for 1H NMR; NT = 16 and NI = 800 for COSY and HSQC; NT = 32 and NI = 1600.

### Determination of *in vitro* antifungal activity of smiglaside C and smilaside A

By running a standard curve with the purified compound using the same LC-MS method with incremental injection volume, the concentrations of smiglaside C and smilaside A were determined to be approximately 0.2 mM and 0.1 mM in *F. graminearum*-induced Mo17 seedling roots, respectively. Purified compounds were re-dissolved in dimethyl sulfoxide (DMSO) to ten-fold of their respective *in vivo* concentrations (*i.e.* 2 mM smiglaside C, and 1 mM smilaside A). An *F. graminearum* spore and hyphae suspension was prepared as described above, 80 uL of it was mixed with 100 uL potato dextrose broth and 20 uL of the testing compounds in a 96-well plate. Twenty uL of pure DMSO were included as the negative control for this experiment. Fungal growth was monitored by light absorbance measurement at 405 nanometers at 30-minute intervals for 12 hours at 28 °C. All treatment groups were measured in at least four replicates for statistical comparison.

### QTL mapping of the constitutive content of smiglaside C, smilaside A, and their ratio

For genetic mapping of constitutive metabolite abundance, three seedlings were germinated for each Intermated B73 × Mo17 (IBM) recombinant inbred line (RIL) and five for each of the parental lines, B73 and Mo17. Ten-day-old root tissues from the three seedlings of each RIL were pooled into one sample for LC-MS analysis, whereas the five B73 and Mo17 seedlings were analyzed individually to allow comparison of their constitutive metabolomes. QTL mapping analysis was performed based on published genotype data using the composite interval mapping algorithm implemented in WinQTL Cartographer v2.5 (Wang, 2012).The significance thresholds of these QTL mapping results were determined with five hundred permutations. For the ratio mapping, the smilaside A/smiglaside C ratio was used.

### Maize root transcriptome analysis

To ensure comparability between metabolome and transcriptome datasets, total RNA was extracted with the Promega SV Total RNA Isolation Kit from another aliquot of the same seedling root tissues used for non-targeted metabolite profiling. In total, seedling root RNA samples were obtained for 73 IBM RILs and five replicates each of B73 and Mo17. mRNA sequencing libraries were prepared robotically on a Biomek NXp with manual post-PCR cleanup using the Lexogen QuantSeq 3′ mRNA-Seq Library Prep Kits (Kremling *et al.*, 2018). These libraries were pooled into one lane and sequenced with 90 base pairs single end reads on an Illumina NextSeq 500 with v2 chemistry at the Cornell Biotechnological Resource Center. Raw RNAseq read data were converted to SAM format, and aligned to B73 RefGen_v3 5b+ gene models using the STAR v2.5.1 RNAseq-aligner (Dobin *et al.*, 2013). The raw transcript counts were calculated for each gene model in each sample using the HTseq 0.6.1p2 python module (Anders *et al.*, 2015). Finally, gene models with fewer than 10 raw read counts in any one of the 73 samples were filtered out, and raw transcript count for each gene model was normalized by the total transcript count of each sample.

### 1-aminocyclopropane-1-carboxylic acid (ACC) and 1-methylcyclopropene (1-MCP) Treatment

*Zmacs2-1 Zmacs6* double mutant seedlings, along with the wildtype B73 progenitor, were grown in Turface (Young *et al.*, 2004). When the seedlings emerged approximately 2 cm above the soil line, they were treated with either 100 uL of 50 mM 1-ACC solution or water control for 5 consecutive days.

In a separate set of experiments, *F. graminearum-*inoculated B73 seedlings were transplanted into Turface and kept in 70 L airtight boxes. The bottom of the boxes were filled with water and EthylBloc (0.14% 1-MCP) sachets to reach a final concentration of 5 g/L 1-MCP. Pots containing maize seedlings were elevated from water surface to avoid direct contact with the solution. 1-ACC and water control treatments were conducted using the same setup to ensure comparability with the 1-MCP treated seedlings. The predicted plant growth effects of 1-ACC and 1-MCP treatments were confirmed by measuring seedling heights in each treatment group.

To confirm that *F. graminearum* growth was not directly influenced by 1-ACC and 1-MCP treatment, plugs of fungal hyphae were transferred to the center of potato dextrose agar plates placed in the sealed 70 L boxes used for seedling experiments. For 1-MCP treatment, EthylBloc was used at the same concentration as described above. For 1-ACC treatment, 50 mM 1-ACC stock solution in DMSO were added to melted and cooled potato dextrose agar to a final concentration of 50 µM. The same final concentration of DMSO was included in plates used for controls and 1-MCP treatments. Fungus radial growth was measured after four days.

For both 1-ACC and 1-MCP experiments, seedling roots were harvested for targeted metabolic analysis with the LC-MS method described above. To confirm that the *Zmacs2-1 Zmacs6* double mutant seedlings are producing less ethylene in their root tissues, 1 mg samples of ground frozen root tissues were placed in airtight 8 mL glass vials for ethylene collection for 29 hours. One mL samples were injected into an Agilent Technologies 6850 Network GC system to estimate ethylene content. The ethylene peak was identified and quantified by comparing to a standard of known concentration, and normalized for tissue weight. Fungus-inoculated seedling roots were examined under an Olympus SZX-12 stereo-microscope with LP Green filter cube to compare fungus spread semi-quantitatively before frozen for LC-MS analysis. Another aliquot of these root tissues was used for q-RT-PCR quantification of *F. graminearum*-specific gene expression to estimate fungal growth, as described above.

### Smiglaside C and smilaside A induction by multiple fungal pathogens

Mo17 stem elicitation assays utilized 35-day-old greenhouse grown plants in 1-L pots. Plants in damage-related treatment groups were slit in the center, spanning both sides of the stem, with a surgical scalpel that was pulled 8 to 10 cm upward to create a parallel longitudinal incision. The damage treatments spanned the upper nodes, internodes, and the most basal portion of unexpanded leaves. *Aspergillus flavus, Rhizopus microspores, Fusarium verticilloides*, and *Cochliobolus heterostrophus* fungal spore inoculations were conducted with 100 µL of water per plant at a concentration of 1 ×10^7^ spores mL^-1^ (Ding *et al.*, 2017). Damage plus water alone was used for a mock inoculation. Localized areas of control and treated stem tissues were covered clear plastic packing tape to minimize tissue desiccation, and stem tissues were harvested 4 d later from each individual plant.

Maize stem tissues were ground to a fine powder in liquid nitrogen and weighed out in 50 mg aliquots. Smilaside A and smiglaside C were analyzed as described previously (Ding *et al.*, 2017). Negative ionization [M-H]^-^ mode scans (0.1-atomic mass unit steps, 2.25 cycles s-1) from m/z 100 to 1,000 were acquired. Analyses of smilaside A and smiglaside C peak abundance relied on the native parent [M-H]^-^ ion m/z 777 and m/z 819, and stable average retention times of 12.76 min and 13.94 min, respectively. Both analytes displayed split peaks and for consistency both peaks where integrated and combined for the final analyses.

### Ergosterol assays

For ergosterol measurements, *F. graminearum*-treated seedlings were transplanted into pots of twice-autoclaved TX360 Metro Mix and grown for 3 weeks, at which point roots were harvested into liquid nitrogen and stored at −80°C. Ergosterol was analyzed as described previously (Christensen *et al.*, 2014). with the following modifications: roots were crushed and placed in scintillation vials each with 10 ml of chloroform:methanol (2:1 v/v) (99.8%) followed by incubation in darkness overnight at room temperature. One ml of extract from each vial was syringe-filtered through 0.45 µm cellulose acetate membrane filters, and 50 µl of filtrate was added to 50 µL of 10 µM C13-labeled cholesterol (cholesterol-25,26,27-13C; Sigma) in methanol as internal standard. Ergosterol was quantified using an Ascentis Express C-18 column (3 cm × 2.1 mm, 2.7 µm) connected to an API 3200 LC/MS/MS with atmospheric photochemical ionization. The injection volume was 5 µl and the isocratic mobile phase consisted of acetonitrile at a flow rate of 300 µl/min.

### Data analysis

All *t-*tests were performed with the ttest function implemented in Microsoft Excel. ANOVA and Wilcoxon rank sum tests were performed in R. Linear discriminant analysis was performed with the SAS software.

## Results

### *Fusarium graminearum* infection leads to root growth inhibition and metabolic reconfiguration

Inoculation of maize genotype B73 with *F. graminearum* under controlled growth conditions, significantly reduced seedling root growth (Figure S1a,b). Using this assay, we screened the 26 parental lines of the maize nested association mapping population (McMullen *et al.*, 2009), as well as inbred lines Mo17 and W22. This showed that B73 was among the most susceptible inbred lines, with root growth being reduced by over 50%. In contrast, Mo17 emerged as a potentially *F. graminearum*-resistant line, showing no significant change in root growth after inoculation (Fig. 1 and Fig. S1c). Further research was focused on the B73 *vs.* Mo17 comparison, due to the availability of genetic resources that included recombinant inbred lines (RILs) and near-isogenic lines (NILs) (Lee *et al.*, 2002; Eichten *et al.*, 2011). The higher fungal resistance of Mo17 was confirmed by lower expression of *FgTUB,* a *F. graminearum*-specific tubulin gene, lower levels of deoxynivalenol, a mycotoxin produced by *F. graminearum*, and fewer visible necrotic symptoms in the roots (Fig. 2a-c).

**Figure 1.**
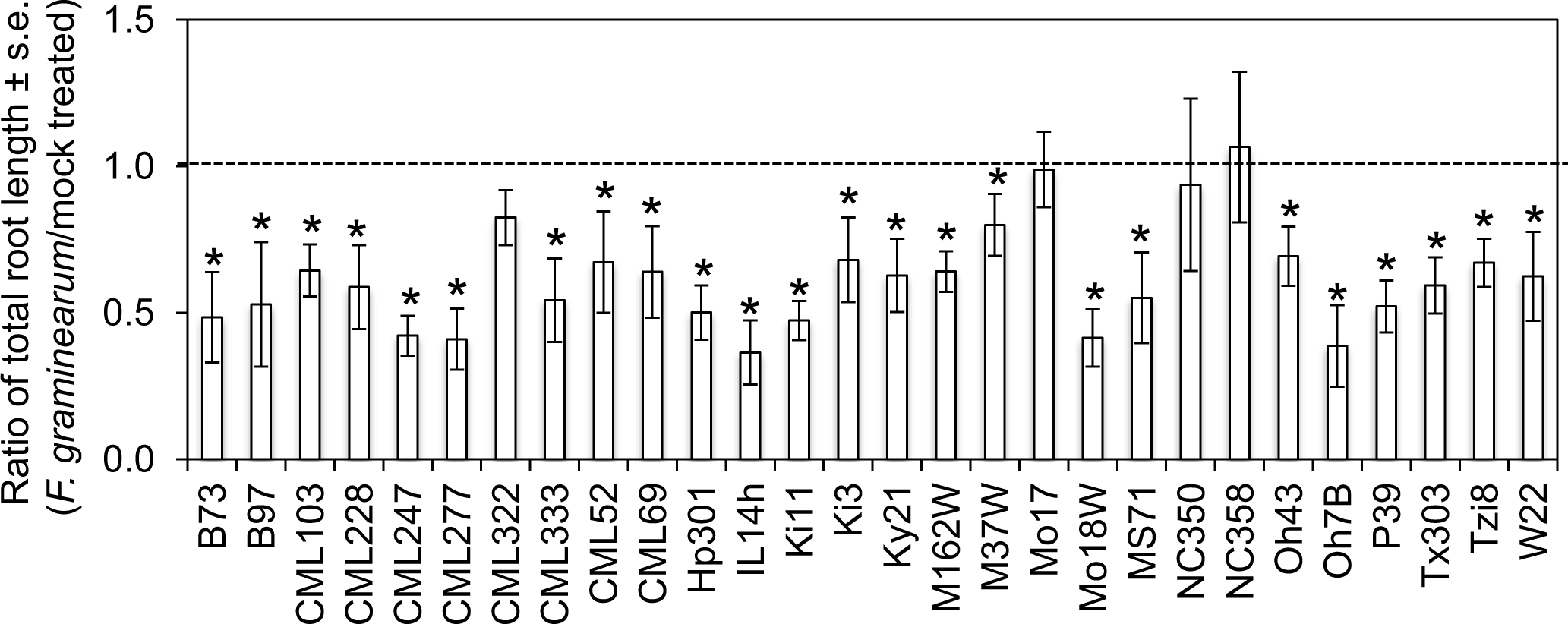
Natural variation in root growth inhibition by *F. graminearum*. The ratio of total root length of *F. graminearum-* and mock-inoculated maize seedlings, measured six days after inoculation, is shown. The dotted line denotes the expected ratio when there is no significant effect of *F. graminearum* infection. At least five mock- and fungus-inoculated individuals are measured for each genotype. *P < 0.05, Student’s *t-*test for significant difference from 1.

**Figure 2.**
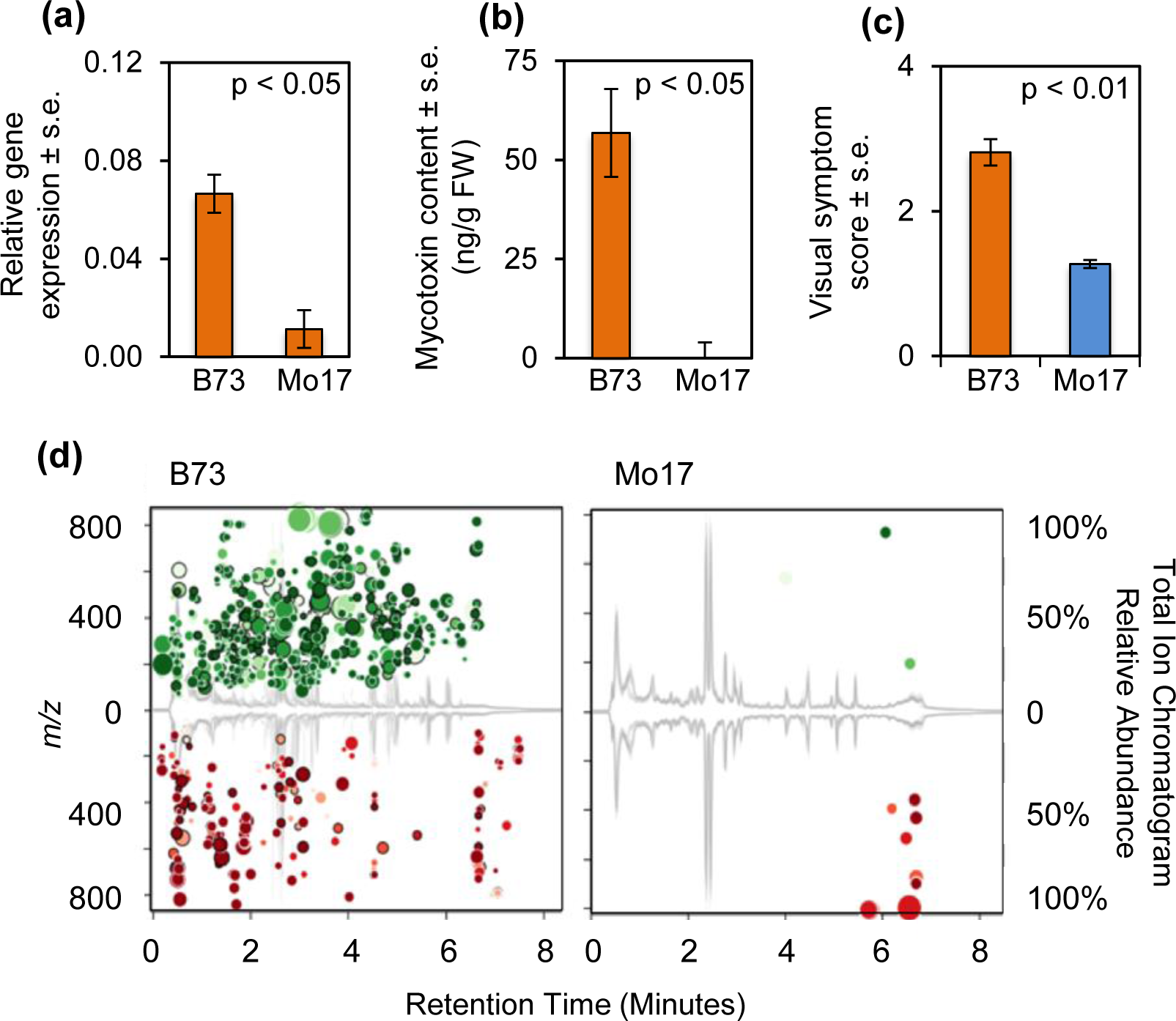
Mo17 is more resistant to *F. graminearum* than B73. Compared to B73, Mo17 seedling roots inoculated with *F. graminearum* demonstrate (a) lower expression of an *F. graminearum-*specific α-tubulin gene, mean +/-s.e. of N = 5, (b) lower content of deoxynivalenol at six-days-post-inoculation, mean +/-s.e. of N = 5, and (c) reduced symptoms on a 0-4 scale, with 0 = no symptoms and 4 = seedling death, mean +/-s.e. of N = 10. *P < 0.05 two-tailed unpaired Student’s *t*-tests for A and B, and paired *t*-test for C. (d) Non-targeted metabolomics of *F. graminearum*-inoculated B73 and Mo17 seedling roots. For either genetic background, each mass feature that was significantly different between treatments (control vs. infected, N = 5; p < 0.05; fold change > 1.5) was plotted as a bubble on the total ion current chromatogram, with its size proportional to the fold change, and darker color representing a more significant change. Mass features induced by *F. graminearum* are shown in the upper half of each plot and *F. graminearum*-suppressed mass features are shown in the lower half.

We hypothesized that the contrasting *F. graminearum* resistance in B73 and Mo17 seedling roots could be attributed to differences in their constitutive and/or inducible biochemical defenses. Therefore, we performed a non-targeted comparative metabolomic analysis of B73 and Mo17 seedling roots, with and without *F. graminearum* inoculation. Consistent with the difference in *F. graminearum*-induced root growth reduction between these two inbred lines, we observed that more than three hundred mass features were significantly altered by *F. graminearum* infection of B73, but only twenty were altered in Mo17 in this experiment (Fig. 2d).

### Acetylated diferuloylsucroses contribute to *F. graminearum* resistance

To identify specific metabolites that could contribute to the contrasting *F. graminearum* resistance levels in B73 and Mo17, a separate non-targeted metabolomic experiment was performed to compare the constitutive metabolomes of B73 and Mo17 seedling roots, as well as mock- and *F. graminearum-*inoculated B73 seedling roots. This identified forty mass features that were both significantly affected by *F. graminearum* in the susceptible B73 seedling roots and constitutively different between B73 and Mo17 seedling roots (Table S1-3). Among these forty mass features, several represented specialized metabolites with known antifungal activity, including benzoxazinoids and phenylpropanoids (Bollina *et al.*, 2010; Ponts *et al.*, 2011; Kazan *et al.*, 2012).

Two mass features with mass-to-charge ratio (*m/z*) of 819.2321 and 777.2221 under negative electron spray ionization (ESI) mode, eluting at 6.11 and 5.61 minutes, respectively, were significantly more abundant after *F. graminearum* infection of both B73 and Mo17 seedling roots. In all samples, the *m/z* 819 metabolite was much more abundant than the *m/z* 777 metabolite (Fig. 3a,b). B73 contains significantly more of the *m/z* 819 metabolite than Mo17, both constitutively and after *F. graminearum* induction (Fig. 3a). In contrast, the *m/z* 777 metabolite was more abundant in Mo17 under both conditions (Fig. 3b).

**Figure 3.**
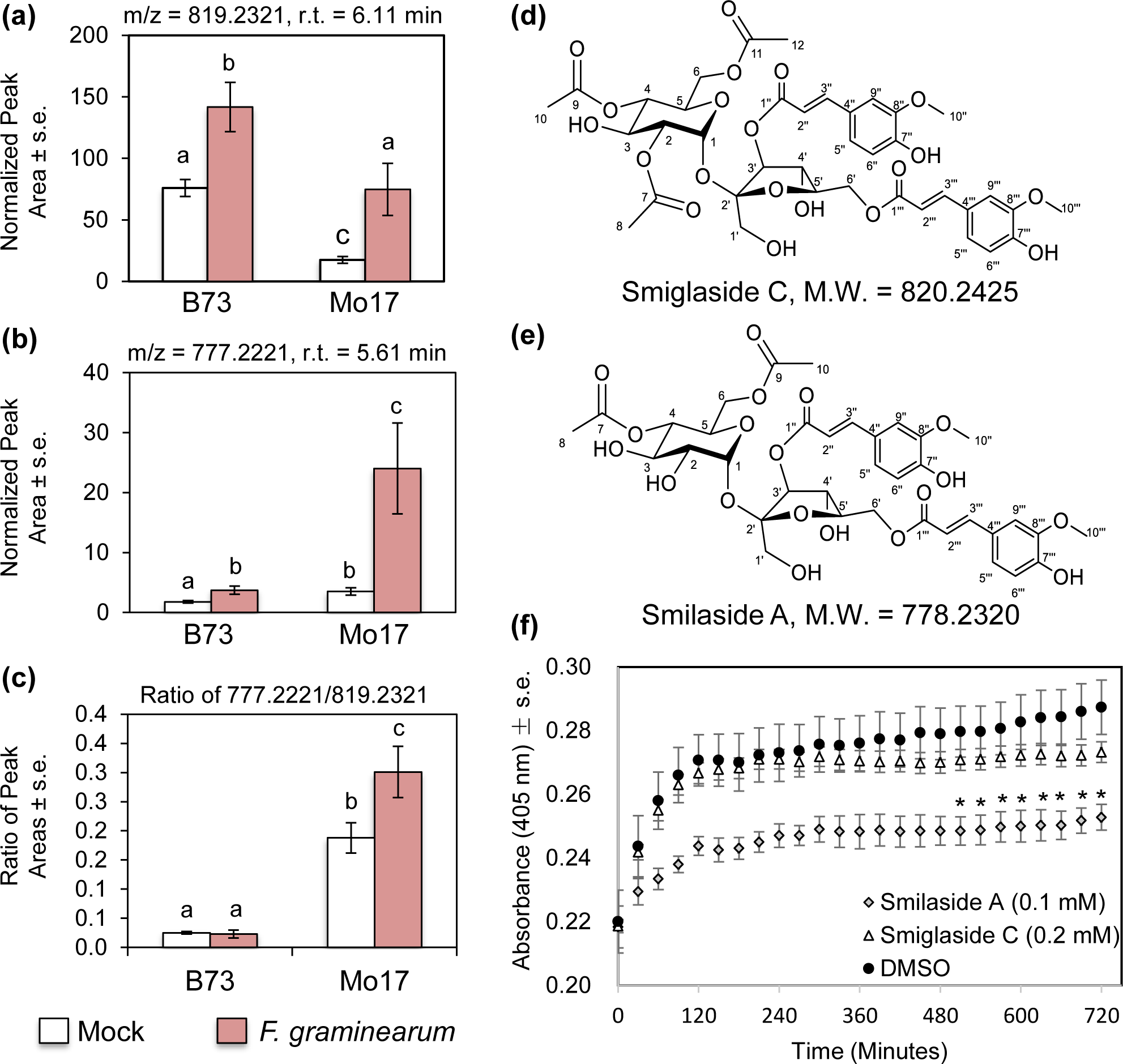
Antifungal metabolites smiglaside C and smilaside A are differentially induced by *F. graminearum* in B73 and Mo17. Abundance of metabolites with (a) *m/z* 819 and (b) *m/z* 777 was measured in negative electron spray ionization mode. (c) Ratio of the peak areas of the two metabolites. Mean +/-s.e. of N = 4. Different letters indicate significant differences, P < 0.05, ANOVA followed by Tukey’s HSD test. Structures of (d) smiglaside C and (e) smilaside A were determined by LC-MS/MS and NMR. (f) Growth of an *F. graminearum* spore/hyphae suspension incubated with smilaside A, smiglaside C, or a dimethylsulfoxide (DMSO) solvent-only control. Fungal growth was monitored by absorbance at 405 nm. Mean +/-s.e. of N = 4, *P < 0.05, Student’s *t*-test relative to the DMSO control at the same time point.

The *m/z* 819 metabolite was identified as 3,6-diferuloyl-2′,3′,6′-triacetylsucrose (Fig. 3d), based on its phenylpropanoid-like UV absorbance profiles (Fig. S2), tandem mass spectrometry (MS/MS; Fig. S2), and nuclear magnetic resonance (NMR) spectroscopy (HSQC, HMBC, and dqfCOSY spectra; Table S4). Based on the MS/MS data and difference in exact mass, the *m/z* 777 metabolite was predicted to have one fewer acetyl group. This was confirmed by one dimensional proton NMR, which showed that the C2′ acetyl group was absent, and the compound was 3,6 diferuloyl-3′,6′-diacetylsucrose (Fig. 3e). These two metabolites were previously identified as smilaside A (3,6 diferuloyl-3′,6′-diacetylsucrose) in *Smilax china* (Kuo *et al.*, 2005; Cho *et al.*, 2015) and smiglaside C (3,6*-*diferuloyl-2′,3′,6′-triacetylsucrose) in *Smilax glabra* (Chen *et al.*, 2000).

In addition to smilaside A and smiglaside C, other maize compounds co-elute with a UV-absorbance peak at 328 nanometers, characteristic of a phenylpropanoid moiety. These include likely structural isomers of smilaside A and smiglaside C, with identical *m/z* ratio and different retention times, as well as possible monoacetylated (*m/z* 735.21) and tetraacetylated (*m/z* 861.24) diferuloylsucroses (Fig. S3). However, the structures of these other maize metabolites were not confirmed, and their functions have not been investigated in this study. Notably, non-acetylated diferuloylsucrose (expected *m/z* ratio = 693.21 in negative ESI mode) was not detected.

The structural resemblance of smilaside A and smiglaside C suggested that they could be the substrate and product of an acetylation reaction, respectively. This acetylation is actively regulated in response to fungal infection, with *F. graminearum* infection inducing a significant increase in the smilaside A/smiglaside C ratio only in the resistant Mo17 seedlings, but not in the susceptible B73 ones (Fig. 3c). This induced response suggested that smiglaside C and/or smilaside A play a role in maize biochemical defense against *F. graminearum*. *In vitro* fungal growth inhibition assays were conducted in liquid suspension culture using smiglaside C and smilaside A concentrations similar to those found in maize seedlings (Fig. S4). Although it was tested at a lower concentration, smilaside A showed a more significant inhibition of *F. graminearum* growth *in vitro* than smiglaside C (Fig. 3f). This was consistent with our earlier observation that the *F. graminearum*-resistant Mo17 seedlings have a higher constitutive smilaside A content, and further accumulated this compound upon fungus attack compared to the susceptible B73 seedlings (Fig. 3a-c).

### Genetic mapping of smiglaside C and smilaside A abundance identifies *ETHYLENE INSENSITIVE 2* as a candidate regulator

The constitutive difference in smilaside A and smiglaside C abundance between B73 and Mo17 seedling roots allowed us to investigate the genetic control of this natural variation. Composite interval mapping with seedling roots of 83 recombinant inbred lines (RILs) from the intermated B73 x Mo17 (IBM) population (Lee *et al.*, 2002; Wang, 2012), together with the two parental lines, showed that the most significant QTL for both metabolites is located at the same position on chromosome 3 (Fig. 4a). In version 2 of the B73 reference genome (Refgen v2), this locus covers approximately 2.3 million base pairs, containing 90 annotated gene models (Schnable *et al.*, 2009; Anders *et al.*, 2015). Interestingly, the two QTL have opposite effects, with the B73 allele promoting constitutive smiglaside C abundance and reducing smilaside A abundance (Fig. 4b,c). Since smilaside A and smiglaside C are likely the substrate-product pair of an acetylation reaction, we hypothesize the mapped QTL regulates the efficiency of this reaction. We identified the same locus when mapping the smilaside A/smiglaside C ratio as a quantitative trait, with the B73 allele reducing the smilaside A/smiglaside C ratio (Fig. S5).

**Figure 4.**
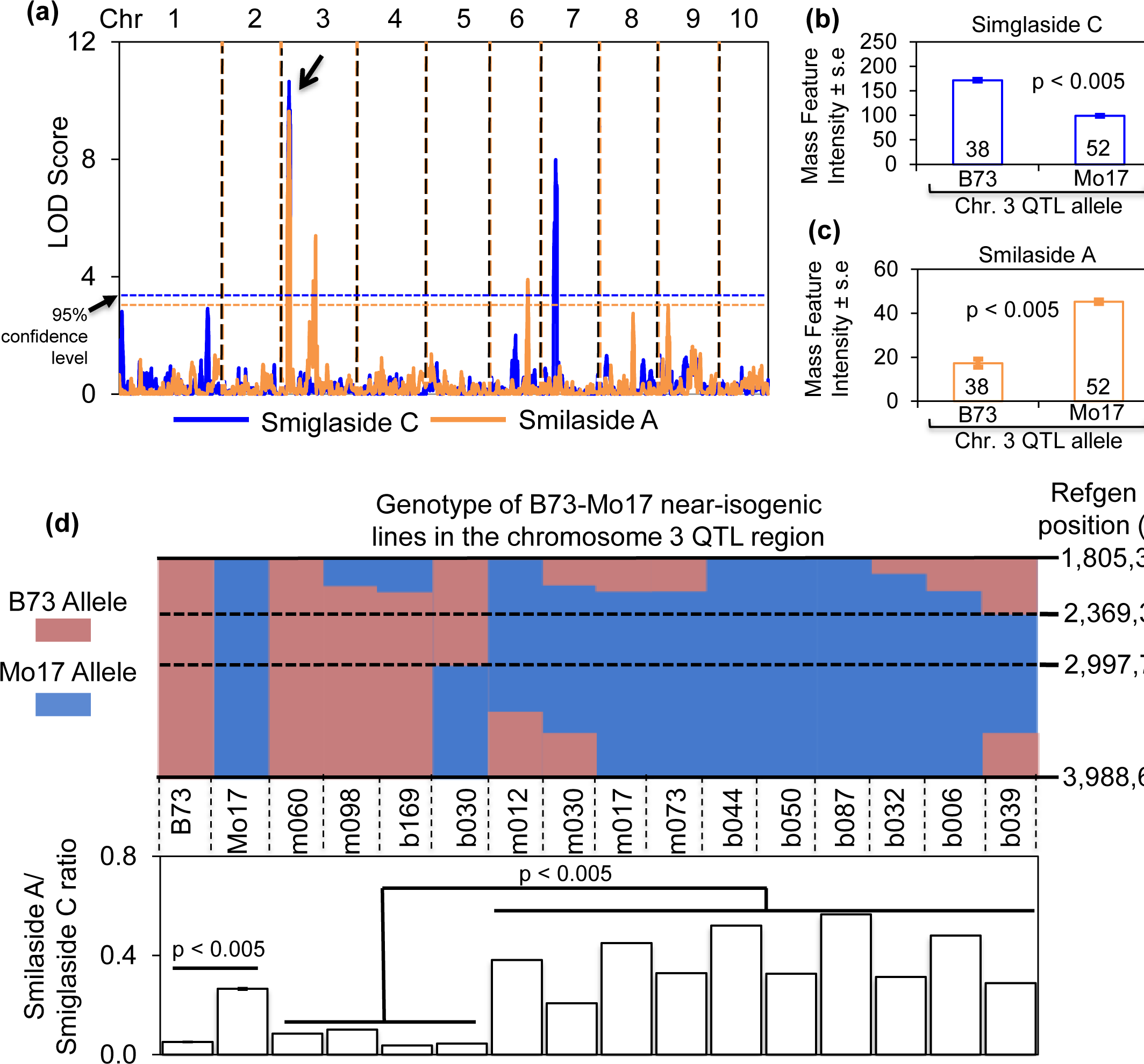
Smiglaside C and smilaside A share a QTL on chromosome 3 with opposite effects. (a) Composite interval mapping of smiglaside C (blue) and smilaside A (orange) in the seedling roots of B73 × Mo17 recombinant inbred lines (RILs) identified a significant QTL for both traits on chromosome 3 (indicated by an arrow). RILs with the B73 QTL allele on chromosome 3 have more smiglaside C (b) and less smilaside A (c) than those with the Mo17 allele. Numbers in bars are sample sizes. P values were determined with two-tailed Student’s *t-* tests. (d) The smilaside A/smiglaside C ratio was calculated for B73, Mo17, and near-isogenic lines. The genetic background of the NILs is indicated by the initial letter of the line name, *e.g.* m060 has a Mo17 genetic background and b169 has a B73 genetic background. The smilaside A/smiglaside C ration is higher in Mo17 than in B73, mean +/-s.e. of N = 3, Student’s *t*-test. NILs carrying the Mo17 allele at the Chromosome 3 QTL have a higher smilaside A/smiglaside C ratio than those with the B73 allele (Students *t-*test), irrespective of the overall genetic background.

To further confirm the role of the QTL in regulating the relative abundance of smiglaside C and smilaside A, we quantified the metabolites in B73-Mo17 near-isogenic lines (NILs) with reciprocal introgressions at this locus (Eichten *et al.*, 2011). Consistent with the RIL results, the smilaside A/smiglaside C ratio showed clear co-segregation with the genetic markers at the chromosome 3 QTL, with the NILs carrying the B73 allele having a lower ratio, irrespective of their genetic background (Fig. 4d). A significant difference between NILs carrying either allele was also observed for smilaside A but not smiglaside C (p = 0.075; Fig. S6). Furthermore, due to additional recombination breakpoints and denser genetic marker data available for the NILs, we narrowed down the QTL region to about 630 kbp, containing 22 predicted gene models.

Natural variation in metabolic traits is often caused by *cis* polymorphisms in metabolic enzyme-encoding genes (Meihls *et al.*, 2013; Yan *et al.*, 2015; Handrick *et al.*, 2016). However, we found no predicted acetyltransferase genes within our QTL interval. Moreover, by plotting the distribution of smiglaside C and smilaside A across the IBM RILs, we found an overall positive correlation between these two metabolites (Fig. 5a), which contradicted the hypothesis of a substrate/product relationship mediated by a polymorphism in a hypothetical acetyltransferase. Closer investigation of the smiglaside C-smilaside A distribution plot revealed that the 83 RILs can be divided by linear discriminant analysis into a B73-like group with lower smilaside A/smiglaside C ratios, and a Mo17-like group with higher ratios (Fig. 5a). We hypothesized that this phenotypic difference could be attributed to transcriptional regulation, which would differ between the IBM RILs belonging to either phenotypic group. We therefore performed whole transcriptome profiling on the seedling root samples that were used for metabolite quantification. This whole-genome analysis showed that the genes with the most significant differential expression between the two phenotypic groups are located in the identified QTL region on chromosome 3 (Fig. 5b,c; Table S5). Specifically, the gene showing the most significantly different expression was a positive regulator of ethylene signaling in maize, *ZmEIN2* (*ETHYLENE INSENSITIVE* 2; GRMZM2G068217), which was expressed at a higher level in the seedling roots of RILs with a B73-like abundance of smiglaside C and smilaside A (Fig. 5d). This leads to the hypothesis that *ZmEIN2*, and hence ethylene signaling, is a negative regulator of smilaside A/smiglaside C ratio.

**Figure 5.**
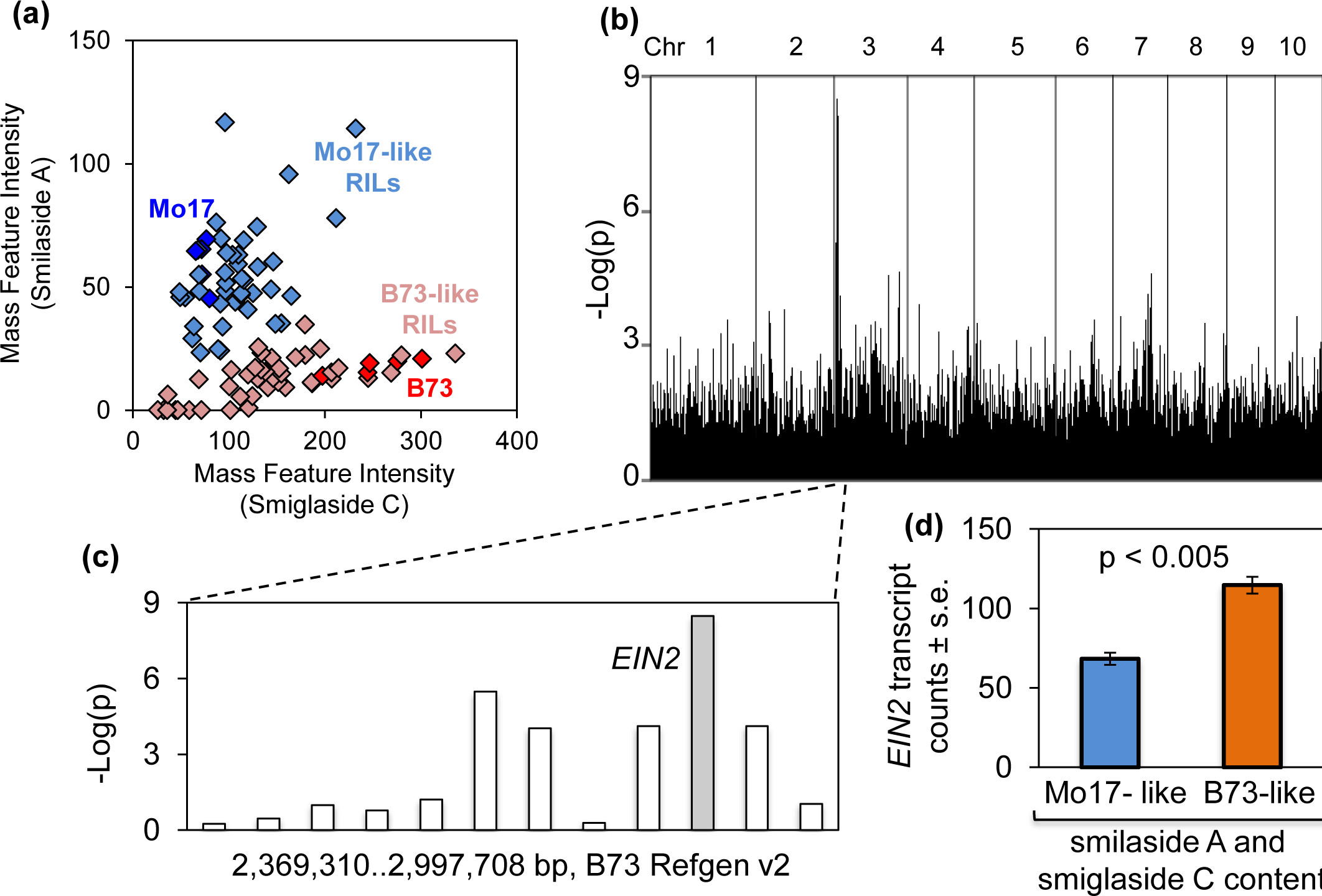
*ETHYLENE INSENSITIVE 2* is differentially expressed in B73 × Mo17 recombinant inbred lines (RILs) with contrasting smilaside A/smiglaside C ratios. (a) B73 × Mo17 RILs can be divided into two groups based on their constitutive smilaside A and smiglaside C content. One replicate each of 83 B73 × Mo17 RILs and five replicates of the B73 and Mo17 parental lines were plotted based on the constitutive content of smilaside A and smiglaside C in their seedling roots. The parental lines are indicated with dark blue (Mo17) and dark red (B73). The RILs were determined to be Mo17-like (light blue) or B73-like (light red) in their smilaside A and smiglaside C content using the linear discriminant analysis. The level of significance in differential expression between Mo17- and B73-like inbred lines, measured by - log(p) from Student’s *t*-tests, is plotted for each root-expressed transcript in the chromosomal order across the whole genome (b) and within the QTL region on chromosome 3 (c). (d) Expression of *EIN2* is significantly higher in the seedling roots of RILs with a B73-like smilaside A and smiglaside C content than in ones with a Mo17-like content. Mean +/- s.e., P value is from a two-tailed Student’s *t-*test.

### Acetylated feruloylsucroses accumulation and resistance against *F. graminearum* are regulated by ethylene

No *ZmEIN2* mutation is available in public maize transposon insertion collections. Instead, we manipulated ethylene response *in vivo* with a gaseous competitive inhibitor, 1-methylcyclopropene (1-MCP), and a biochemical precursor of ethylene production in plant, 1-aminocyclopropane-1-carboxylic acid (1-ACC). Effects of these treatments were confirmed by contrasting seedling growth rate in these groups (Fig. S7). Consistent with the genetic mapping results, 1-MCP treatment (and hence lower *ZmEIN2* availability) led to hyper-accumulation of smilaside A and an elevated smilaside A/smiglaside C ratio (Fig. 6a,c). Unexpectedly, smiglaside C abundance was also significantly increased by 1-MCP treatment (Fig. 6b). Both smilaside A and smiglaside C were induced by 1-ACC treatment, but their ratio was not significantly affected (Fig. 6a-c).

**Figure 6.**
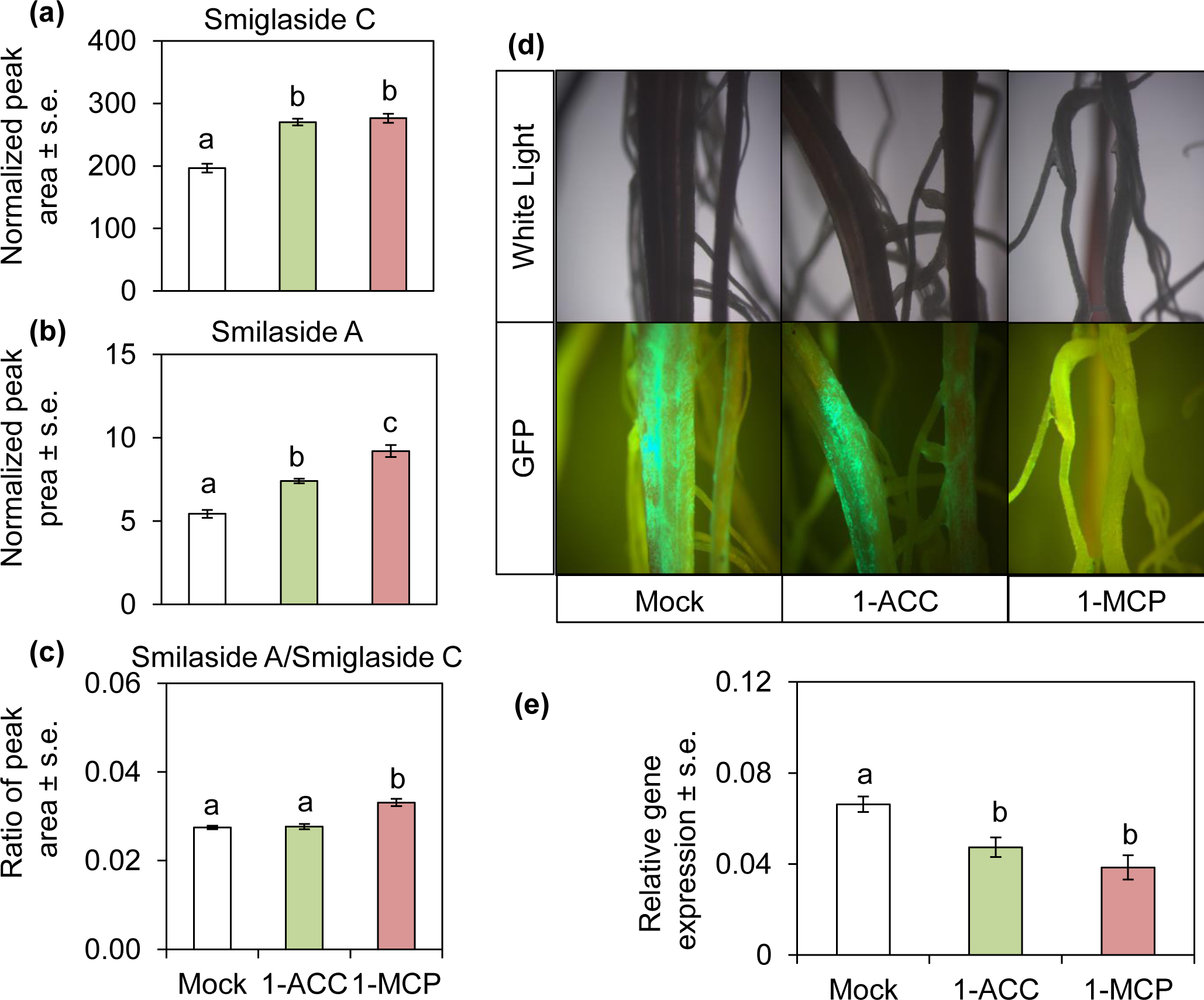
Exogenous 1-methylcyclopropane (1-MCP) treatment promotes maize seedling root resistance against *F. graminearum*. B73 maize seedling roots inoculated with *F. graminearum* were treated with 1-aminocyclopropane-1-carboxylate (1-ACC), 1-MCP, or water as a control. The abundance of (a) smiglaside C, (b) smilaside A, and (c) the ratio of the two metabolites was calculated from peak area under negative electron spray ionization mode. Mean +/-s.e. of N = 5, different letters indicate significant difference, P < 0.05, ANOVA followed by Tukey’s HSD test. (d) B73 maize seedling roots inoculated with *F. graminearum-*GFP and treated with 1-ACC, 1-MCP, or mock treatment for ten days were examined with white light and fluorescence microscopy. More GFP marker expression from *F. graminearum* was observed on root surface of mock- and 1-ACC treated seedling roots than on 1-MCP-treated roots. (e) Fungal growth was quantified by qRT-PCR using *F. graminearum-*specific primers, relative to expression measurement of a maize housekeeping gene. Mean +/- s.e. of N = 8, different letters indicate significant differences, P < 0.05, ANOVA followed by Tukey’s HSD test.

To further investigate how smilaside A and smiglaside C are regulated by ethylene production, we measured their abundance in the seedling roots of the *Zmacs2-1 Zmacs6* ethylene biosynthetic mutant (Young *et al.*, 2004), which has *Mutator* transposon insertions in two 1-ACC synthase genes in the B73 genetic background. Consistent with prior measurement of lower leaf ethylene content (Young *et al.*, 2004), this double mutant had a lower root ethylene concentration than wildtype (Fig. S8). Metabolite abundance was measured with and without exogenous 1-ACC, an ethylene biosynthesis intermediate that is downstream of the two mutated *ZmACS* genes. Constitutively, there were significantly lower amounts of both smilaside A and smiglaside C in the roots of *Zmacs2-1 Zmacs6* compared to wildtype B73. After 1-ACC treatment, both metabolites were increased in wildtype B73, and were restored to wildtype levels in *Zmacs2-1 Zmacs6* (Fig. 7a,b). In *Zmacs2-1 Zmacs6*, the smilaside A/smiglaside C ratio was significantly lower than in wildtype B73. However, consistent with results from the previous experiment, 1-ACC treatment did not affect this ratio in either genetic background (Fig. 7c).

**Figure 7.**
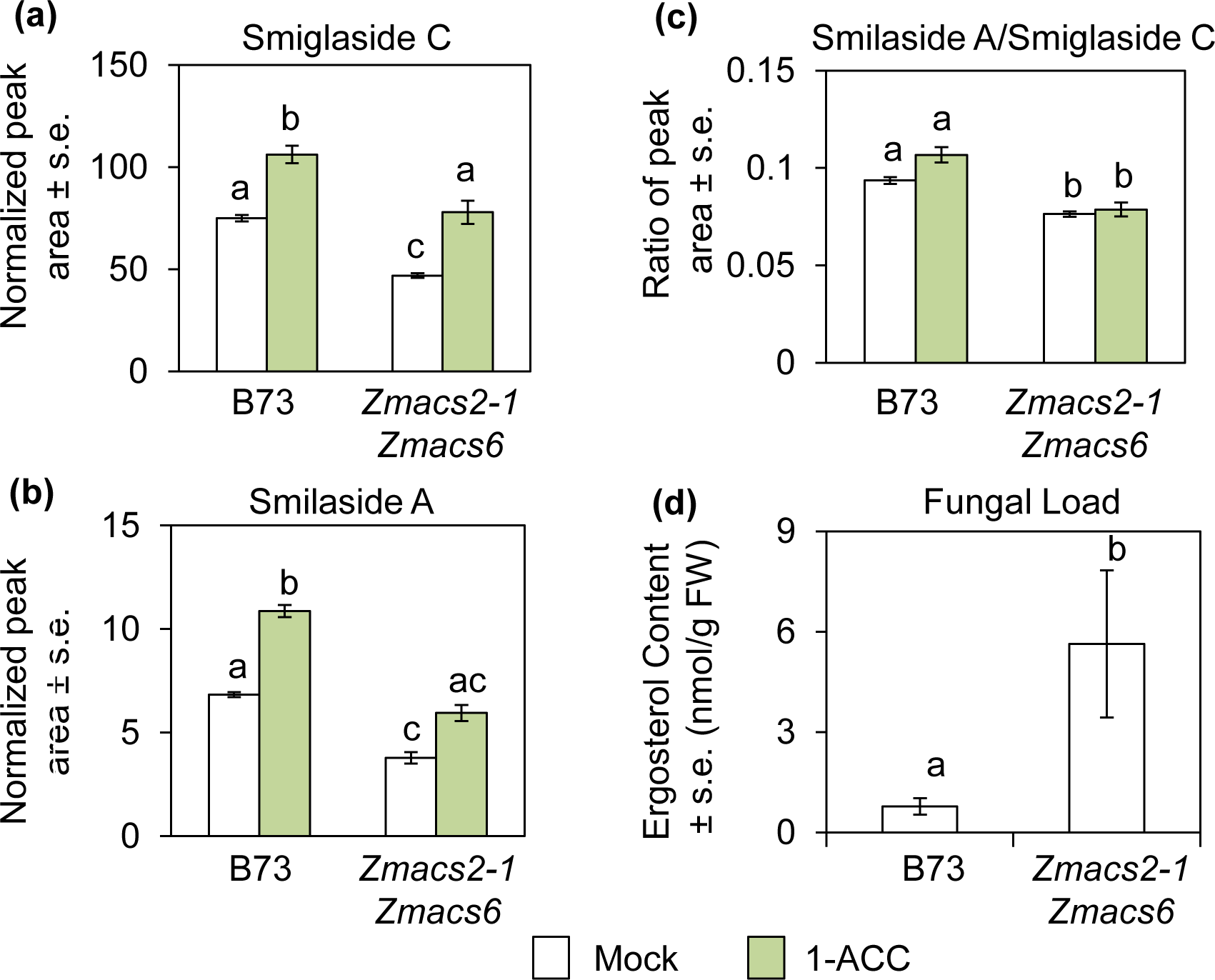
Ethylene biosynthesis is required for acetylated feruloylsucrose accumulation and resistance against *F. graminearum*. The abundance of (a) smiglaside C and (b) smilaside A was estimated by peak area at their respective *m/z* ratio under negative electron spray ionization mode. (c) The ratio of smilaside A and smiglaside C peak areas. Mean +/- s.e., different letters indicate significant differences, P < 0.05, Two-way ANOVA followed by Tukey’s HSD test. (d) Fungal load was estimated by ergosterol content, normalized by root tissue fresh weight. Mean +/- s.e., different letters indicate significant differences, P < 0.05, One-way ANOVA.

To investigate how ethylene production and sensitivity can affect maize seedling defense against *F. graminearum,* we compared *F. graminearum-*inoculated maize seedling roots treated with 1-ACC, 1-MCP, or water control. Although we observed extensive fungal hyphae of GFP-transformed *F. graminearum* on the root surface of both mock and 1-ACC treated seedlings, hyphae were almost completely absent from 1-MCP treated roots (Fig. 6d). In support of the microscopic observations, we found significantly lower *FgTUB* expression in 1-MCP treated seedling roots (Fig. 6e). Neither 1-ACC nor 1-MCP affected growth of *F. graminearum* on agar plates (Fig. S9), indicating that there is not direct toxic effect. Comparing infected seedling roots of *Zmacs2-1 Zmacs6* and wildtype B73, we found significantly higher *F. graminearum* growth on the mutant line (as measured by ergosterol accumulation; Fig. 7d).

### Maize diferuloylsucroses are induced by multiple fungal pathogens

A mass feature likely representing smiglaside C was identified as a maize acyl sugar that was induced after infection with *Colletotrichum graminicola* (anthracnose leaf blight), though without structural confirmation (Balmer *et al.*, 2013). To determine whether induced production of smilaside A and smiglaside C was a more general maize response to fungal infection, we inoculated seedlings with four additional fungal pathogens, *Aspergillus flavus, Rhizopus microspores, Fusarium verticilloides*, and *Cochliobolus heterostrophus*. Whereas smilaside A was only induced by *F. verticilloides* and *C. heterostrophus*, all four pathogens significantly induced the accumulation of smiglaside C (Fig. 8). As in the case of *F. graminearum* infection (Fig. 3a-c), smiglaside C was induced to a greater extent than smilaside A. Therefore, induced accumulation of smiglaside C may be a general response of maize to infection by fungal pathogens.

**Figure 8.**
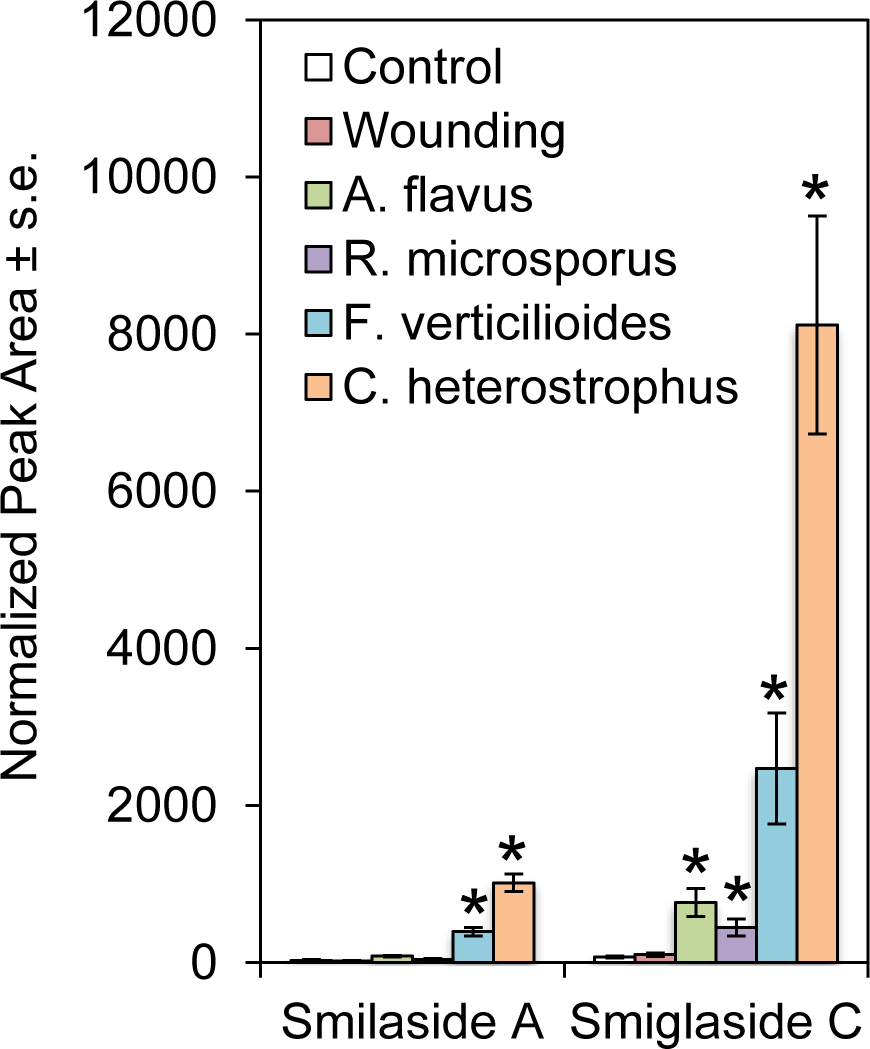
Smilaside A and smiglaside C are induced by fungal pathogen infection. Smilaside A and smiglaside C content were measured in 35-day-old Mo17 stem tissues four days after physical wounding or inoculation with *Aspergillus flavus, Rhizopus microspores, Fusarium verticilloides*, or *Cochliobolus heterostrophus*. Mean +/- s.e. of N = 4. *P < 0.05 relative to the uninfected control, pairwise Wilcoxon rank sum tests with Benjamini and Hochberg correction for multiple comparisons.

## Discussion

In addition to maize, acetylated feruloylsucroses may have defensive properties in other plant species. These compounds were first identified in the rhizomes of *S. china* and *S. glabra*, which are used in traditional Chinese medicine (Chen *et al.*, 2000; Kuo *et al.*, 2005). Similar phenylpropanoid sucrose esters, with different numbers and types of phenylpropanoid groups attached, were later found in various Liliaceae and Polygonaceae species. Crude plant extracts containing these compounds, and in some cases purified compounds, have shown anticancer and antioxidant activities *in vitro* (Zhu *et al.*, 2006; Ono *et al.*, 2007; Yan *et al.*, 2008; Zhang *et al.*, 2008; Kim *et al.*, 2010). Building on these promising *in vitro* bioactivities, organic synthesis routes to produce natural phenylpropanoid sucrose esters and structural analogs have been developed with moderate yield and selectivity (Panda *et al.*, 2012a; Panda *et al.*, 2012b). More recently, acetylated feruloylsucroses and other phenylpropanoid sucrose esters were found in rice (Chen *et al.*, 2014; Cho *et al.*, 2015).

Our discovery of smilaside A and smiglaside C in maize will facilitate the *in planta* investigation of phenylpropanoid sucrose ester function and metabolism. Due to the lack of suitable maize mutants, we could not experimentally prove a causal relationship between the genetic polymorphism in *ZmEIN2* and natural variation in constitutive smiglaside C and smilaside A abundance. Instead, we demonstrated that ethylene biosynthesis was required for the production of both compounds *in vivo*, whereas ethylene sensitivity fine-tuned their relative abundance (Fig. 6 and 7). Interestingly, fungal load after *F. graminearum* inoculation was negatively correlated with smilaside A/smiglaside C ratio across three natural variation and manipulative experiments in this study (Fig. 3, 6, and 7), suggesting that this ratio, rather than the absolute abundance of these two compounds, may be more important for resistance against *F. graminearum*.

*Fusarium graminearum* infection is known to induce ethylene biosynthetic and responsive genes in both maize seedling roots and *Brachypodium distachyon* spikes (Pasquet *et al.*, 2014). In wheat, different comparative transcriptomic studies have reached opposite conclusions regarding the role of ethylene signaling in responses to *F. graminearum* infection (Li & Yen, 2008; Ding *et al.*, 2011; Xiao *et al.*, 2013). However, *F. graminearum* resistance could be manipulated by manipulating ethylene signaling in both wheat and barley leaves (Chen *et al.*, 2009). In the current study, we observed that maize seedlings with lower ethylene sensitivity, either due to genetically-encoded polymorphism in *ZmEIN2* expression or artificial treatment with a competitive inhibitor, are more resistant to *F. graminearum* and accumulated the more bioactive smilaside A (Fig. 3 and 6). However, exogenous ethylene supplementation in the form of 1-ACC did not promote *F. graminearum* susceptibility in maize seedling roots. This inconsistency could arise from differences in tissue type, developmental stage, or the treatment regime. Together, these results indicate that, above a certain minimal ethylene concentration, ethylene sensitivity negatively regulates the efficiency of biochemical defense, leading to contrasting fungal resistance levels.

In our *in vitro* assay, we observed that the diacetylated smilaside A caused greater fungal growth inhibition than the triacetylated smiglaside C (Fig. 3f). This is perhaps surprising, because phenylpropanoid sucrose esters with higher degrees of acetylation have generally shown stronger *in vitro* bioactivities, though these two specific compounds have not been compared previously (Panda *et al.*, 2012b; Cho *et al.*, 2015). Our observations may be explained by a non-linear relationship between the degrees of acetylation and bioactivity. Different structural isomers also may be relevant for the structure-activity-relationship. Other than the putative tetraacetylated diferuloylsucrose, all acetylated diferuloylsucroses detected in our LC-MS analyses showed signs of multiple structural isomers, though there was usually a predominant one (Fig. S2). How these different isomers could differ in their bioactivities is another question that requires further investigation.

Since acetylated feruloylsucroses can be induced by various fungal pathogens in maize, it would be interesting to assess their crop protection value *in vivo* under more relevant field conditions. Simple phenylpropanoids can contribute to disease resistance not only through their direct antimicrobial activities, but also by playing a critical role in physical fortification of plant cell walls (Nicholson & Hammerschmidt, 1992). The importance of cell walls as a physical barrier against *F. graminearum* and other fungal pathogens is further highlighted by the prevalence of genes that are likely to encode cell-wall-degrading enzymes in the genomes of fungal phytopathogens (Cuomo *et al.*, 2007; Kubicek *et al.*, 2014). Should acetylated feruloylsucroses also contribute to the physical strength of plant cell walls, such effects would not be evident in *in vitro* assays.

Although our study has not revealed any enzyme-encoding genes that are directly involved in the biosynthesis of the metabolites of interest, the chemical structures of smilaside A and smiglaside C give us hints about their possible biosynthetic pathway. Specifically, we hypothesize that distinct but related hydroxycinnamoyl transferases are responsible for the esterification of feruloyl-CoA and the free hydroxyl groups on the sucrose molecule, analogous to the phenylpropanoid quinic acid esterification reactions. In the B73 genome, there are 13 predicted hydroxycinnamoyl transferase-encoding genes, mostly with unconfirmed activity and substrates (Schnable *et al.*, 2009). These predicted gene models are candidates for elucidation of the biosynthetic pathway of acetylated feruloylsucroses in maize. Compared to the feruloyl esterification enzymes, the identities of the acetyltransferases that catalyze the acetylation on the glucose ring are less clear. We speculate that these enzymes probably belong to the diverse BAHD acyltransferase family, similar to the acylsugar acyltransferases found in tomatoes (Kim *et al.*, 2012; Schilmiller *et al.*, 2012). Our results lead to the prediction that one or more of these acyltransferases would be positively regulated by ethylene signaling in maize seedling roots. The exact order of feruloyl esterification and acetylation on the sucrose molecule also remains unclear. In rice, non-acetylated 3,6 diferuloylsucrose is detected at a very low level in bulk root extract, suggesting that the acetylation occurs after feruloyl esterification (Cho *et al.*, 2015). However, we did not detect the same compound in our microliter-scale LC-MS analyses of maize roots

Finally, this study demonstrates the feasibility of combining metabolomics, transcriptomics and quantitative genetics methods to elucidate regulation of previously unknown antifungal metabolites in maize. An expansion of this integrative approach to a larger number of maize inbred lines in a genome-wide association study likely will identify both additional metabolites and genes involved in their metabolism. Such genes will be useful in future breeding efforts to enhance the pathogen resistance during maize seedling establishment.

## Acknowledgements

This research was funded by US National Science Foundation awards to 1339237 GJ and 1139329 to GJ and EAS, and US National Science Foundation Graduate Research Fellowship Program award DGE-1650441 to KAK. We thank Chong Huang for help with the linear discrimination analysis, Eric Craft and Jon Schaff for help with the RootReader 2D platform, Francis Trail and Robert Procter for sharing the *F. graminearum* ZTE strain, Navid Mohaved for assistance with mass spectrometry, Julia Vrebalov for assistance with ethylene measurement, Eli Borrego for his assistance in ergosterol analyses, and Annett Richter and Melkamu Woldemariam for helpful discussions.

## Author Contributions

S.Z. and G.J. designed experiments, performed or assisted in all experiments, and wrote the manuscript. K.K. and E.B. prepared the QuantSeq 3′ mRNA-Seq libraries. K.Y.Z. and F.S. purified metabolites and performed NMR spectroscopy analyses. J.S.Ba. and D.K. assisted in the root imaging experiment and the *Zmacs2-1 Zmacs6* mutant analysis, respectively. E.A.S., Y.D., M.V.K., and J.S.Be. conducted fungal infections and *Zmacs2-1 Zmacs6* experiments.

## List of Supporting Information

**Figure S1.** Natural variation in F. *graminearum*-induced root growth reduction in maize seedlings.

**Figure S2.** Predicted feruloylsucrose compounds have characteristic phenylpropanoid-like ultraviolet absorbance profiles.

**Figure S3.** Tandem MS spectra of putative acetylated feruloylsucroses.

**Figure S4.** Standard curve of smiglaside C

**Figure S5.** QTL mapping of the smilaside A/smiglaside C ratio identifies the same locus on Chromosome 3.

**Figure S6.** Constitutive abundance of smilaside A and smiglaside C vary across B73-Mo17 isogenic lines.

**Figure S7.** Endogenous ethylene production in root is depleted in the *Zmacs2-1 Zmacs6* double mutant maize seedlings.

**Figure S8.** 1-Aminocyclopropane-1-carboxylic acid and 1-monocyclopropene treatment has opposite effects on maize seedling height.

**Figure S9.** 1-Aminocyclopropane-1-carboxylic acid and 1-monocyclopropene are not directly toxic to F. *graminearum*.

**Table S1.** List of mass features significantly different between B73 and Mo17 seedling roots constitutively.

**Table S2.** List of mass features significantly different between mock- and *F. graminearum-* inoculated B73 seedling roots.

**Table S3.** List of mass features both significantly different between B73 and Mo17 seedling roots constitutively, and responsive to *F. graminearum* inoculation in B73.

**Table S4.** Chemical shifts and major coupling constants from NMR spectroscopy analysis of smiglaside C.

**Table S5.** Normalized and filtered transcript count of gene models located in the QTL interval across intermated B73-Mo17 recombinant inbred lines and the two parental inbred lines.

---

**Figure S1.**
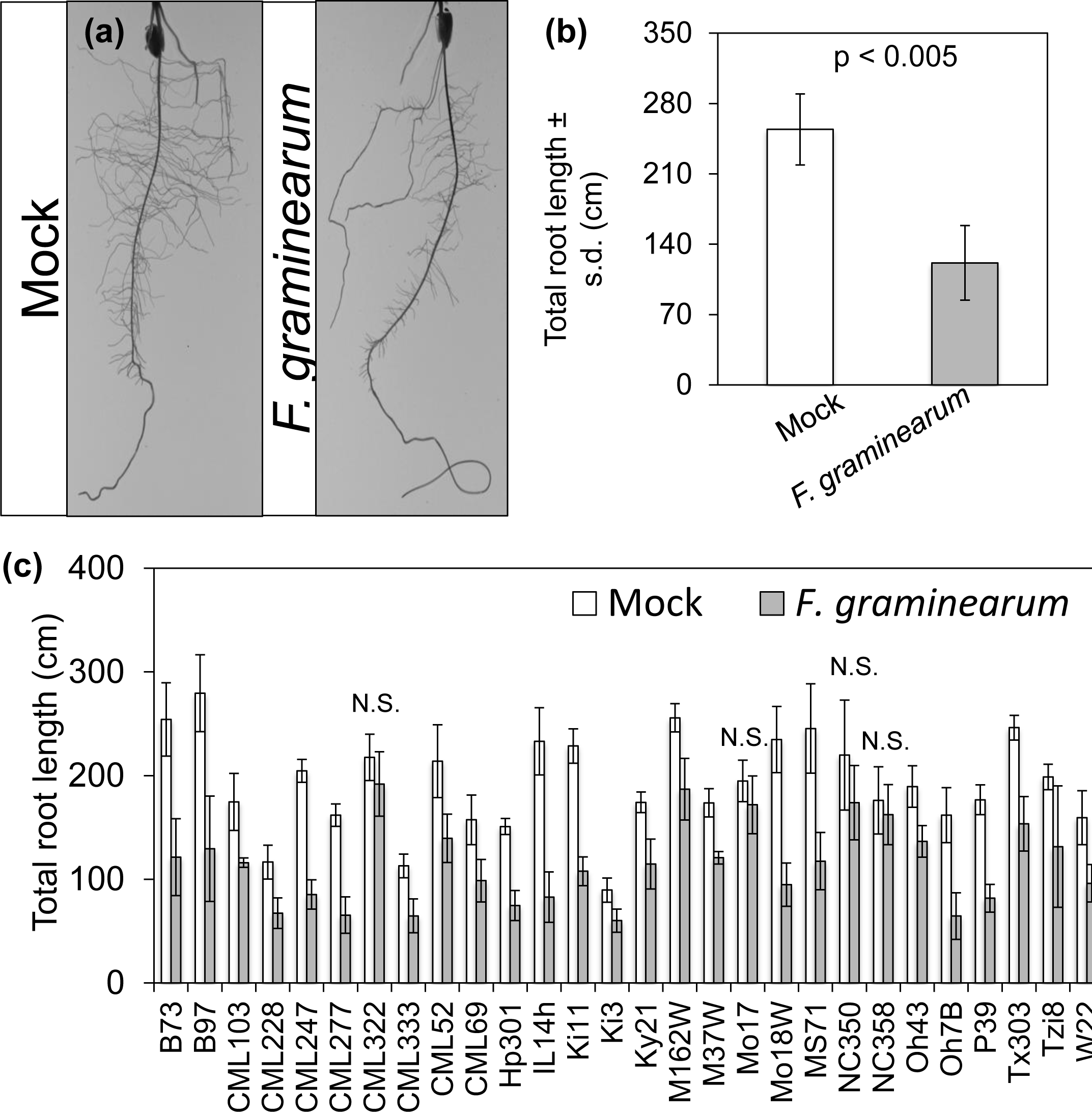
Natural variation in *F. graminearum-*induced root growth inhibition. (A) Representative photos of ten-day-old B73 maize seedling roots mock-inoculated or inoculated with a *F. graminearum* spore suspension. Photos were taken at six days post-inoculation. Similar photos were used for total root length measurement. (B) Eight replicates of either mock- or *F. graminearum-*inoculated B73 seedlings were measured to identify differences between the treatment groups, P < 0.05, Student’s *t-*test. (C) The same assay was expanded to include the 26 parental lines of the nested association mapping population, along with inbred lines Mo17 and W22, to identify natural variation in *F. graminearum*-induced maize seedling root growth reduction. All but four lines (marked by N.S.) showed significant reduction in root growth six days after *F. graminearum* inoculation (P < 0.05, two-tailed Student’s *t-*test). Root growth ratios based on these data are shown in Figure 1.

**Figure S2.**
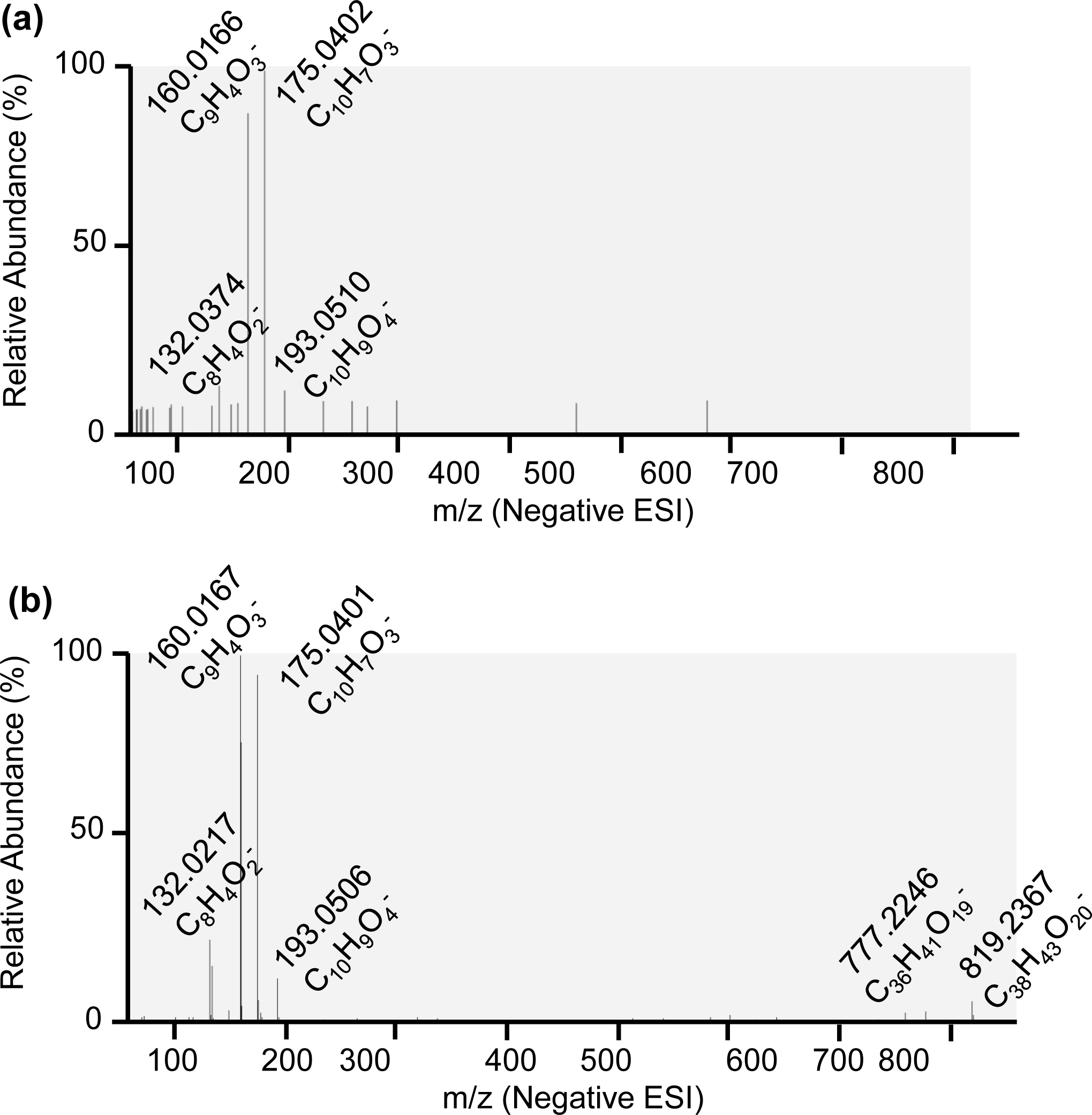
Tandem MS spectra of predicted acetylated feruloylsucroses. The tandem MS spectra of the two putative acetylated feruloylsucroses were generated from mock-inoculated Mo17 and B73 seedling root extracts. Scan filters were set to an *m/z* ratio range around their expected parentalion. Major daughter ions are labeled with their exact *m/z* value and most probable predicted molecular formula.

**Figure S3.**
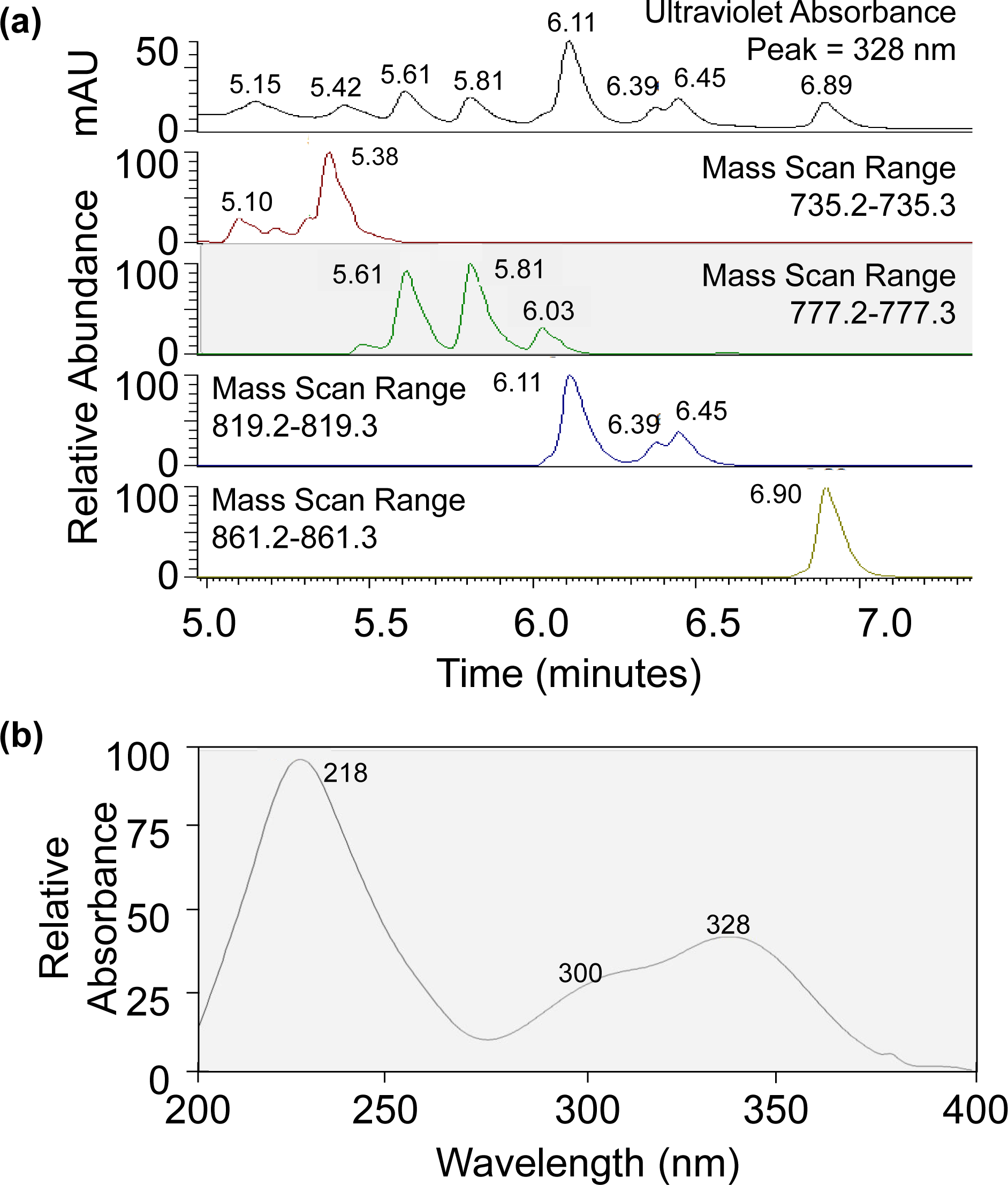
Predicted feruloylsucrose compounds have characteristic phenylpropanoid-like ultraviolet absorbance profiles. Feruloylsucrose compounds, as predicted by their exact molecular mass, co-elute with UV absorbance peaks with characteristic phenylpropanoid-like absorbance profiles. UV absorbance data collected from a photo diode array detector are filtered to show only peak absorbance at 328 nm, and mass spectrometry data are filtered for the expected molecular masses of predicted feruloyl acetylsucroses (a). Each peak is labeled with its retention time. A characteristic phenylpropanoid-like UV absorbance profile is shown, with peaks at 218 nm and 328 nm, and a shoulder at 300 nm (b).

**Figure S4.**
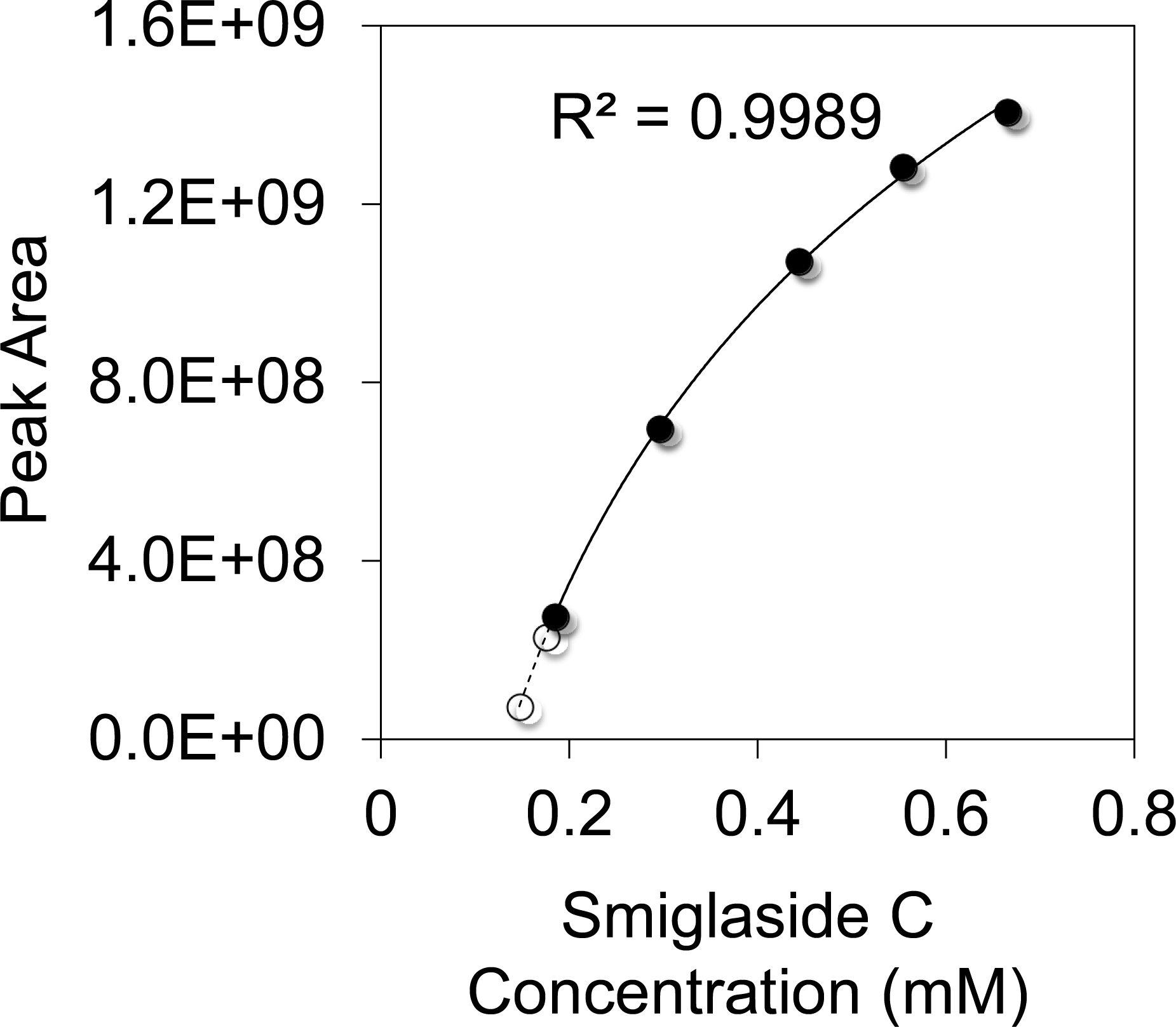
Standard curve of smiglaside C. Calculated smiglaside C concentrations from five different injection volumes of the same purified compound solution are plotted with their corresponding mass feature peak areas in filled marks. A logarithmic curve was fitted to these data points with a calculated regression co-efficient (R2). The same curve is extended to calculate the *in vivo* concentrations of smiglaside C (0.1750 mM) and smilaside A (0.1470 mM) based on MS peak areas in an *F. graminearum-*induced Mo17 seedling root extract. These two peak areas from biological samples and their inferred concentrations are shown in empty circles.

**Figure S5.**
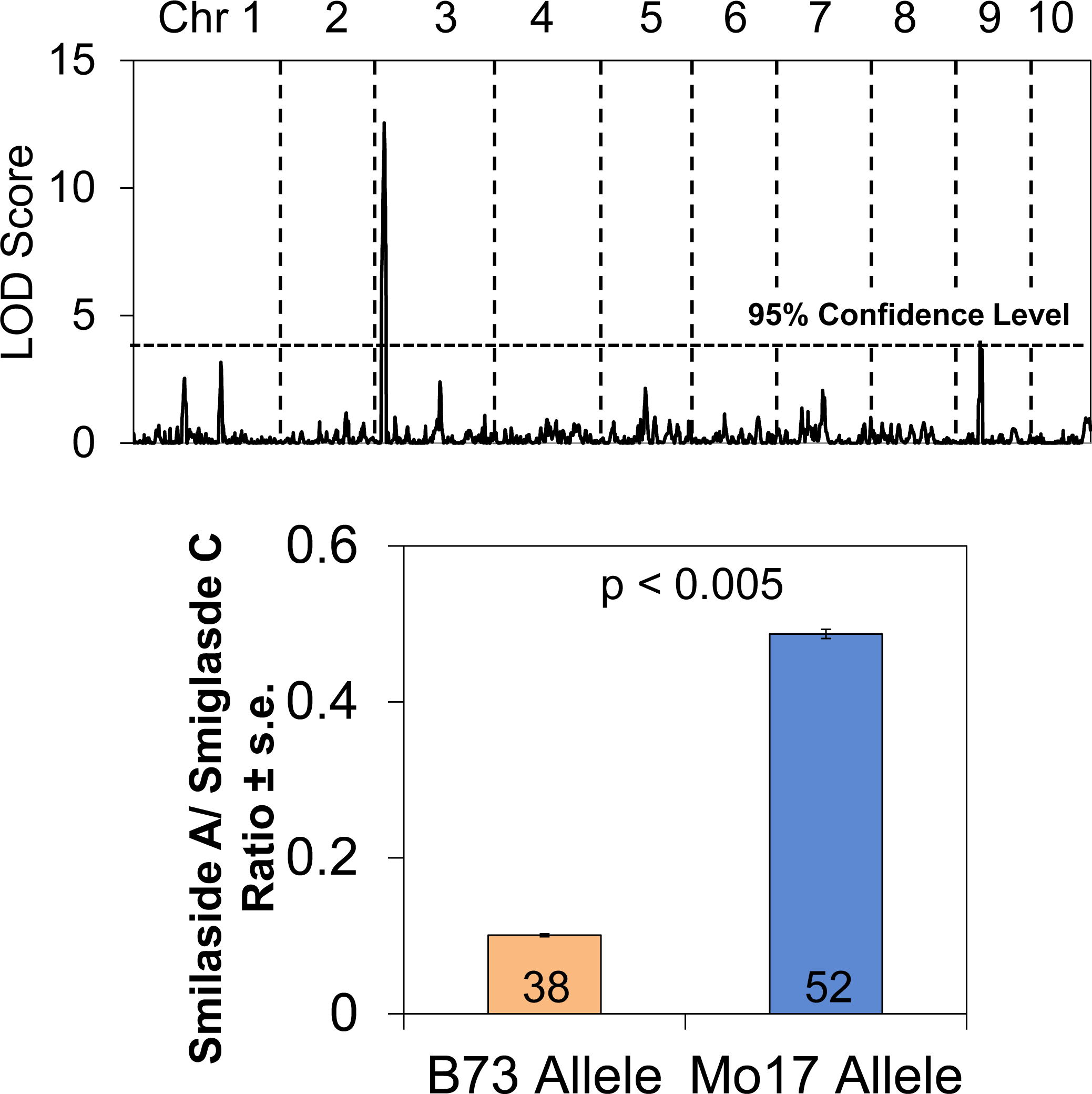
QTL mapping of the smilaside A/smiglaside C ratio identifies a locus on Chromosome 3. The smilaside A/smiglaside C ratio in the seedling roots of 83 B73 × Mo17 recombinant inbred lines (RILs) was used for composite interval mapping. The calculated log of odds (LOD) score (top panel) of each locus is plotted across the ten maize chromosomes. The 95% confidence interval of the LOD scores was calculated with 500 permutations. RILs and parental lines carrying the Mo17 allele at the most significant QTL on chromosome 3 have a significantly higher smilaside A/smiglaside C ratio than those carrying the B73 allele (lower panel). P-values were calculated using Student’s t-test, and the number of RILs in either group is indicated each column.

**Figure S6.**
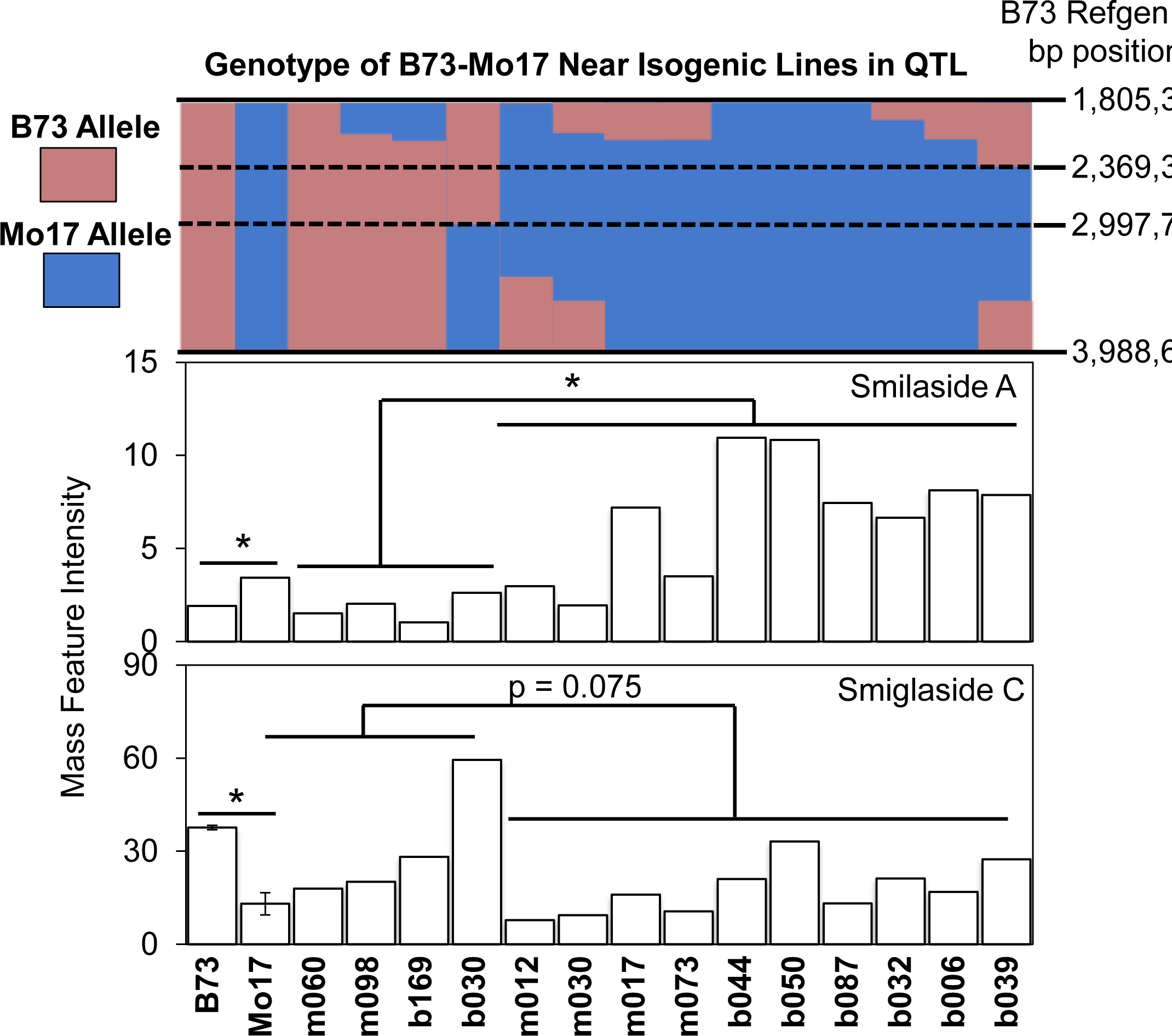
Constitutive abundance of smilaside A and smiglaside C varies across B73 × Mo17 near isogenic lines (NILs). The genetic region mapped in the top panel is determined by the 95% confidence interval of a chromosome 3 QTL for smilaside A and smiglaside C abundance. The genetic map of this region and the allelic state of each near isogenic line are reproduced based on data published in Eichten et al., 2011. The genetic background of the NILs is indicated by the initial letter of the line name, *e.g.* m060 has a Mo17 genetic background and b169 has a B73 genetic background Constitutive abundance of smilaside A and smiglaside C were estimated from the mass feature intensity measured in roots of B73, Mo17, and NILs. Mean +/- s.e. of N = 5, *P < 0.05, Student’s *t-*test.

**Figure S7.**
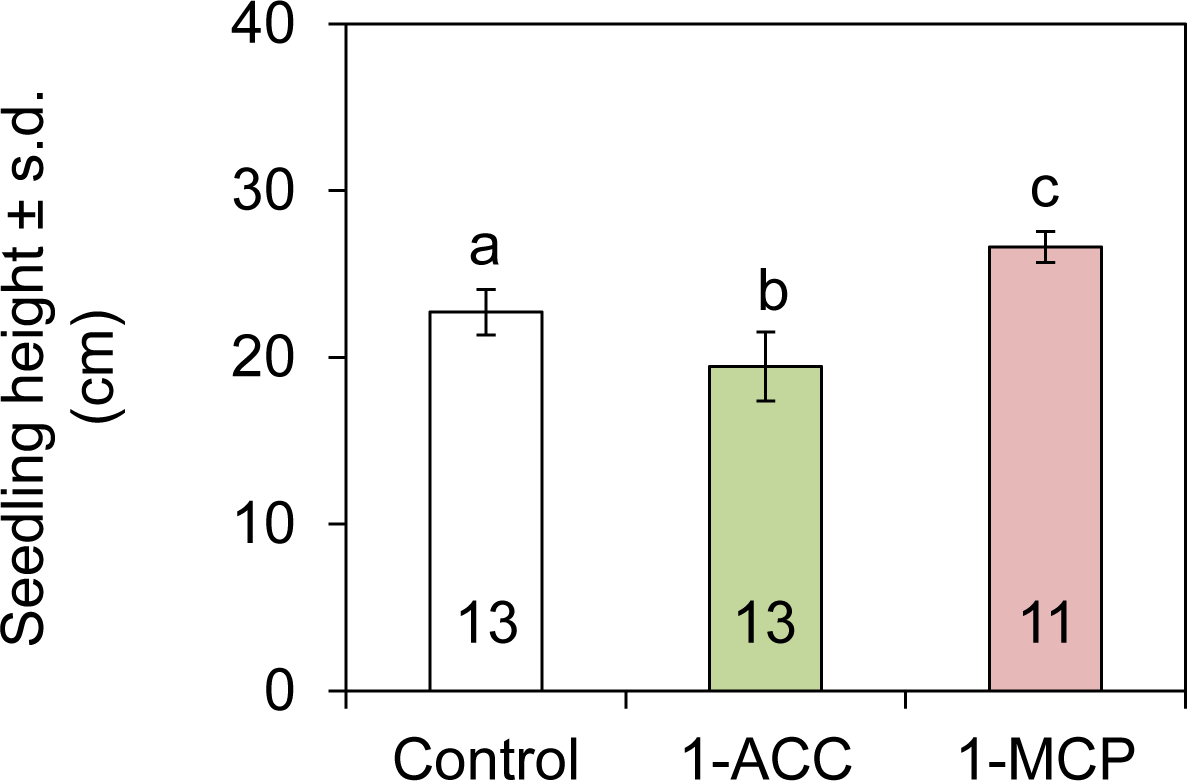
1-Aminocyclopropane-1-carboxylic acid and 1-monocyclopropene treatments have opposite effects on maize seedling height. Seedling heights between each pair of treatment groups are compared to control with one-way ANOVA and Tukey’s HSD test, and significant differences (P < 0.05) are indicated by different letters. The number of biological replicates is denoted by numbers in each bar.

**Figure S8.**
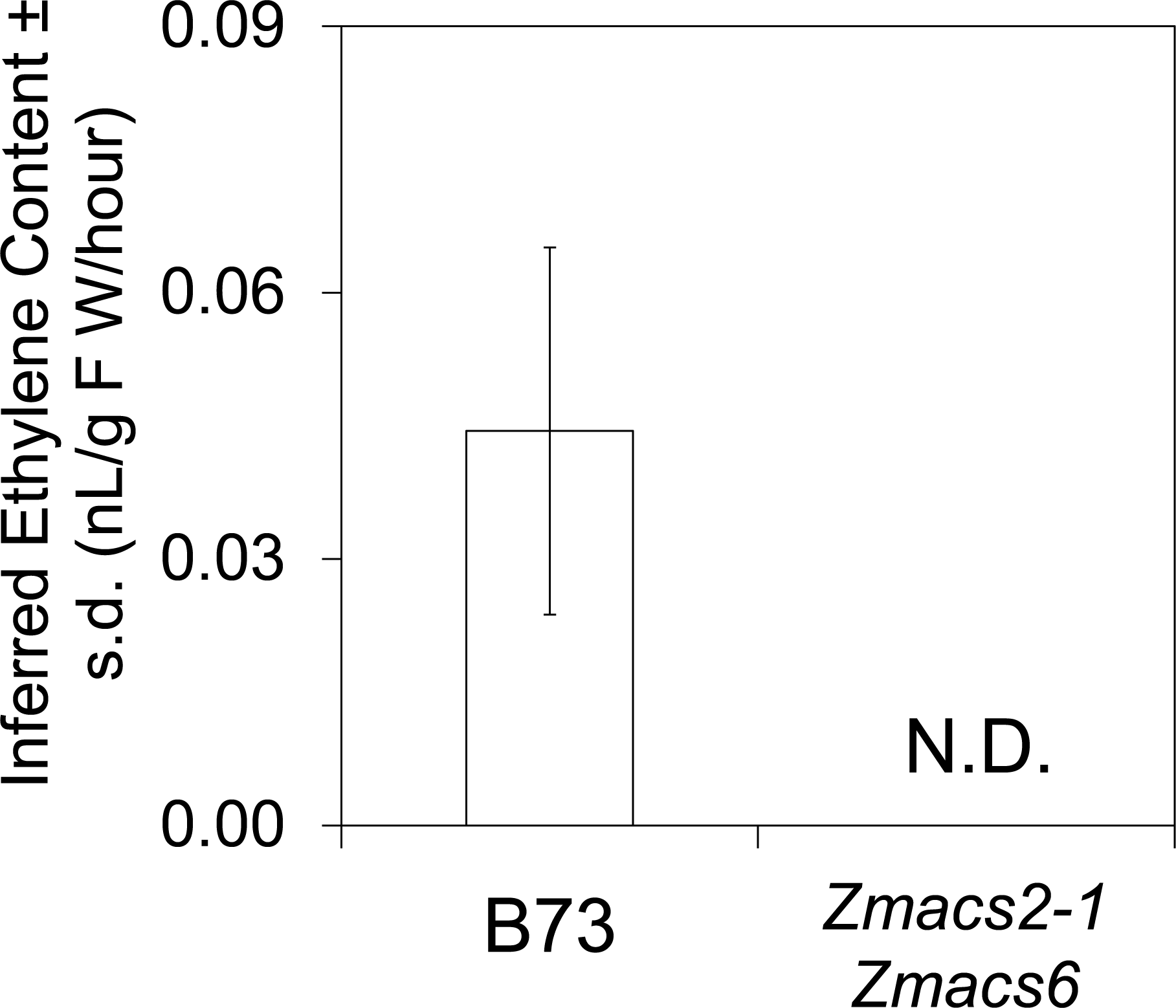
Endogenous ethylene production in root is depleted in the *Zmacs2-1 Zmacs6* double mutant maize seedlings. Ethylene produced by root tissues was collected in sealed glass vials for 29 hours and analyzed by gas chromatography. Absolute ethylene content is inferred from peak area measurement from a flame ionization detector. Peaks corresponding to ethylene were not detected (N.D.) in the *Zmacs2-1 Zmacs6* samples. Mean+/- s.e. of N = 6.

**Figure S9.**
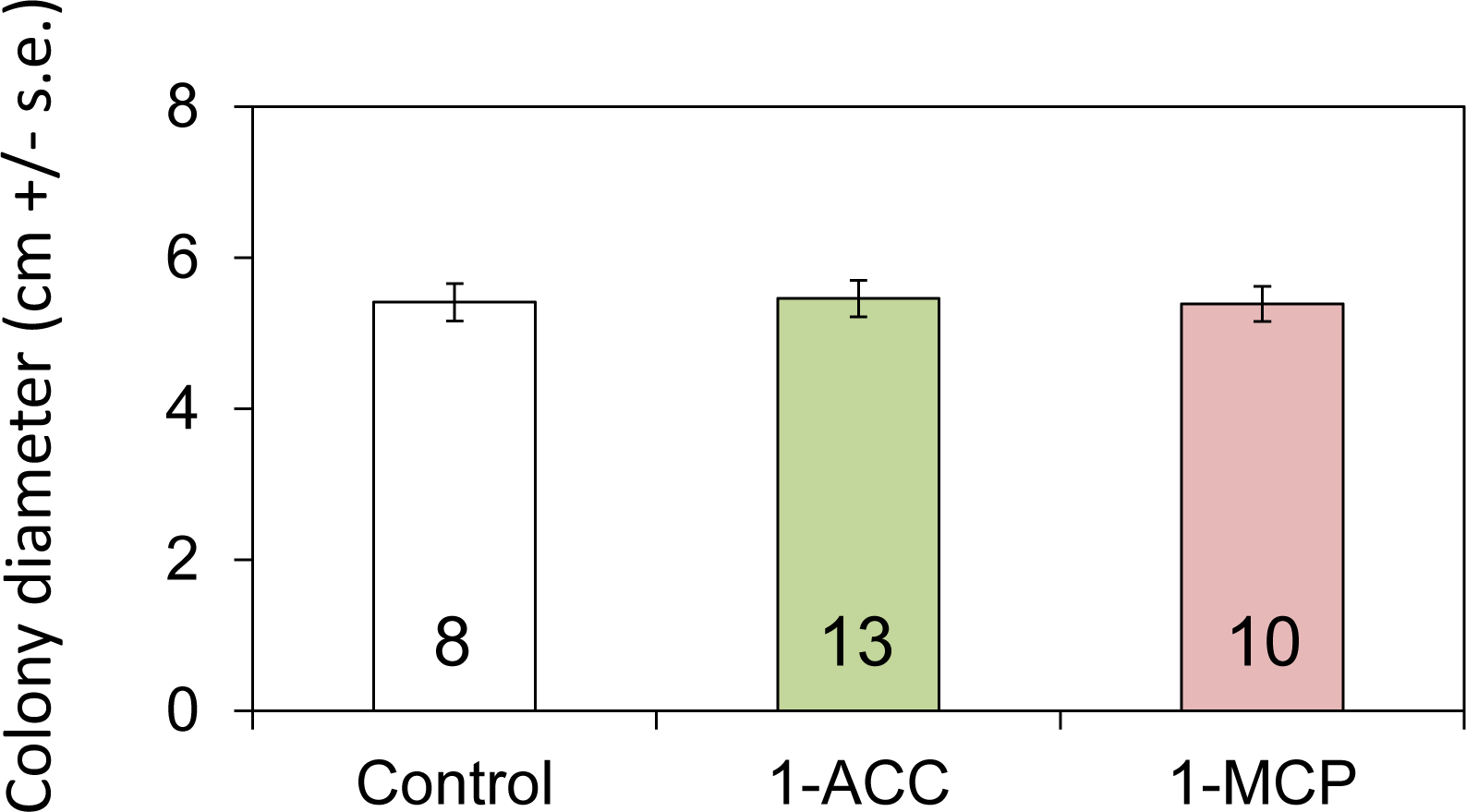
1-Aminocyclopropane-1-carboxylic acid and 1-monocyclopropene are not directly toxic to *F. graminearum.* Fungi were inoculated into the center of potato dextrose agar plates, enclosed in airtight boxes, and colony diameter was measured after five days. 1-ACC was added to the agar at 50 µMconcentration. 1-MCP was released as a volatile using EthylBloc at 5 g/L. Mean +/- s.e., numbers in bars indicate sample sizes, no significant differences P > 0.05, two-tailed Student’s *t-*tests.

## References

Ali ML, Taylor JH, Jie L, Sun G, William M, Kasha KJ, Reid LM, Pauls KP. 2005. Molecular mapping of qtls for resistance to gibberella ear rot, in corn, caused by fusarium graminearum. Genome 48: 521–533.

Anders S, Pyl PT, Huber W. 2015. Htseq: A python framework to work with high-throughput sequencing data. Bioinformatics 31: 166–169.

Balmer D, de Papajewski DV, Planchamp C, Glauser G, Mauch-Mani B. 2013. Induced resistance in maize is based on organ-specific defence responses.Plant Journal 74: 213–225.

Benson JM, Poland JA, Benson BM, Stromberg EL, Nelson RJ. 2015. Resistance to gray leaf spot of maize: Genetic architecture and mechanisms elucidated through nested association mapping and near-isogenic line analysis.PLoS Genet 11.

Benton HP, Want EJ, Ebbels TMD. 2010. Correction of mass calibration gaps in liquid chromatography-mass spectrometry metabolomics data.Bioinformatics 26: 2488–2489.

Bollina V, Kumaraswamy GK, Kushalappa AC, Choo TM, Dion Y, Rioux S, Faubert D, Hamzehzarghani H. 2010. Mass spectrometry-based metabolomics application to identify quantitative resistance-related metabolites in barley against fusarium head blight. Molecular Plant Pathology 11: 769–782.

Brauner PC, Melchinger AE, Schrag TA, Utz HF, Schipprack W, Kessel B, Ouzunova M, Miedaner T. 2017. Low validation rate of quantitative trait loci for gibberella ear rot resistance in european maize. Theor Appl Genet 130: 175–186.

Buckler ES, Gaut BS, McMullen MD. 2006. Molecular and functional diversity of maize. Curr Opin Plant Biol 9: 172–176.

Chen T, Li JX, Xu Q. 2000. Phenylpropanoid glycosides from smilax glabra. Phytochemistry 53: 1051–1055.

Chen W, Gao Y, Xie W, Gong L, Lu K, Wang W, Li Y, Liu X, Zhang H, Dong H, et al. 2014. Genome-wide association analyses provide genetic and biochemical insights into natural variation in rice metabolism. Nat Genet 46: 714–721.

Chen X, Steed A, Travella S, Keller B, Nicholson P. 2009. Fusarium graminearum exploits ethylene signalling to colonize dicotyledonous and monocotyledonous plants. New Phytologist 182: 975–983.

Cho JG, Cha BJ, Seo WD, Jeong RH, Shrestha S, Kim JY, Kang HC, Baek NI. 2015. Feruloyl sucrose esters from oryza sativa roots and their tyrosinase inhibition activity. Chemistry of Natural Compounds 51: 1094–1098.

Christensen SA, Nemchenko A, Park YS, Borrego E, Huang PC, Schmelz EA, Kunze S, Feussner I, Yalpani N, Meeley R, et al. 2014. The novel monocot-specific 9-lipoxygenase zmlox12 is required to mount an effective jasmonate-mediated defense against fusarium verticillioides in maize. Mol Plant Microbe Interact 27: 1263–1276.

Cuomo CA, Gueldener U, Xu JR, Trail F, Turgeon BG, Di Pietro A, Walton JD, Ma LJ, Baker SE, Rep M, et al. 2007. The fusarium graminearum genome reveals a link between localized polymorphism and pathogen specialization. Science 317: 1400–1402.

Ding LN, Xu HB, Yi HY, Yang LM, Kong ZX, Zhang LX, Xue SL, Jia HY, Ma ZQ. 2011. Resistance to hemi-biotrophic f. Graminearum infection is associated with coordinated and ordered expression of diverse defense signaling pathways. Plos One 6.

Ding Y, Huffaker A, Kollner TG, Weckwerth P, Robert CAM, Spencer JL, Lipka AE, Schmelz EA. 2017. Selinene volatiles are essential precursors for maize defense promoting fungal pathogen resistance. Plant Physiology doi: 10.1104/pp.17.00879. [Epub ahead of print].

Dobin A, Davis CA, Schlesinger F, Drenkow J, Zaleski C, Jha S, Batut P, Chaisson M, Gingeras TR. 2013. Star: Ultrafast universal rna-seq aligner. Bioinformatics 29: 15–21.

Eichten SR, Foerster JM, de Leon N, Kai Y, Yeh CT, Liu S, Jeddeloh JA, Schnable PS, Kaeppler SM, Springer NM. 2011. B73-mo17 near-isogenic lines demonstrate dispersed structural variation in maize. Plant Physiology 156: 1679–1690.

Famoso AN, Clark RT, Shaff JE, Craft E, McCouch SR, Kochian LV. 2010. Development of a novel aluminum tolerance phenotyping platform used for comparisons of cereal aluminum tolerance and investigations into rice aluminum tolerance mechanisms. Plant Physiology 153: 1678–1691.

Guenther JC, Trail F. 2005. The development and differentiation of Gibberella zeae (anamorph: Fusarium graminearum) during colonization of wheat. Mycologia 97: 229–237.

Handrick V, Robert CAM, Ahern KR, Zhou SQ, Machado RAR, Maag D, Glauser G, Fernandez-Penny FE, Chandran JN, Rodgers-Melnik E, et al. 2016. Biosynthesis of 8-o-methylated benzoxazinoid defense compounds in maize. Plant Cell 28: 1682–1700.

Huffaker A, Kaplan F, Vaughan MM, Dafoe NJ, Ni XZ, Rocca JR, Alborn HT, Teal PEA, Schmelz EA. 2011. Novel acidic sesquiterpenoids constitute a dominant class of pathogen-induced phytoalexins in maize. Plant Physiology 156: 2082–2097.

Jiao Y, Peluso P, Shi J, Liang T, Stitzer MC, Wang B, Campbell MS, Stein JC, Wei X, Chin CS, et al. 2017. Improved maize reference genome with single-molecule technologies. Nature 546: 524–527.

Kazan K, Gardiner DM, Manners JM. 2012. On the trail of a cereal killer: Recent advances in fusarium graminearum pathogenomics and host resistance. Molecular Plant Pathology 13: 399–413.

Kebede AZ, Woldemariam T, Reid LM, Harris LJ. 2016. Quantitative trait loci mapping for gibberella ear rot resistance and associated agronomic traits using genotyping-by-sequencing in maize. Theor Appl Genet 129: 17–29.

Kim J, Kang K, Gonzales-Vigil E, Shi F, Jones AD, Barry CS, Last RL. 2012. Striking natural diversity in glandular trichome acylsugar composition is shaped by variation at the acyltransferase2 locus in the wild tomato solanum habrochaites. Plant Physiology 160: 1854–1870.

Kim KH, Chang SW, Lee KR. 2010. Feruloyl sucrose derivatives from bistorta manshuriensis. Canadian Journal of Chemistry-Revue Canadienne De Chimie 88: 519–523.

Kremling KAG, Chen SY, Su MH, Lepak NK, Romay MC, Swarts KL, Lu F, Lorant A, Bradbury PJ, Buckler ES. 2018. Dysregulation of expression correlates with rare-allele burden and fitness loss in maize. Nature 555: 520–523.

Kubicek CP, Starr TL, Glass NL. 2014. Plant cell wall-degrading enzymes and their secretion in plant-pathogenic fungi. Annual Review of Phytopathology 52: 427–451.

Kuhl C, Tautenhahn R, Bottcher C, Larson TR, Neumann S. 2012. Camera: An integrated strategy for compound spectra extraction and annotation of liquid chromatography/mass spectrometry data sets. Analytical Chemistry 84: 283–289.

Kuo YH, Hsu YW, Liaw CC, Lee JK, Huang HC, Kuo LMY. 2005. Cytotoxic phenylpropanoid glycosides from the stems of smilax china. Journal of Natural Products 68: 1475–1478.

Lee M, Sharopova N, Beavis WD, Grant D, Katt M, Blair D, Hallauer A. 2002. Expanding the genetic map of maize with the intermated b73 x mo17 (ibm) population. Plant Mol Biol 48: 453–461.

Li GL, Yen Y. 2008. Jasmonate and ethylene signaling pathway may mediate fusarium head blight resistance in wheat. Crop Science 48: 1888–1896.

Lou YR, Bor M, Yan J, Preuss AS, Jander G. 2016. Arabidopsis nata1 acetylates putrescine and decreases defense-related hydrogen peroxide accumulation. Plant Physiology 171: 1443–1455.

McMullen MD, Kresovich S, Villeda HS, Bradbury P, Li H, Sun Q, Flint-Garcia S, Thornsberry J, Acharya C, Bottoms C, et al. 2009. Genetic properties of the maize nested association mapping population. Science 325: 737–740.

Meihls LN, Handrick V, Glauser G, Barbier H, Kaur H, Haribal MM, Lipka AE, Gershenzon J, Buckler ES, Erb M, et al. 2013. Natural variation in maize aphid resistance is associated with 2,4-dihydroxy-7-methoxy-1,4-benzoxazin-3-one glucoside methyltransferase activity. Plant Cell 25: 2341–2355.

Mideros SX, Windham GL, Williams WP, Nelson RJ. 2012. Tissue-specific components of resistance to aspergillus ear rot of maize. Phytopathology 102: 787–793.

Mueller D 2016a. Corn disease loss estimates from the united states and ontario, canada - 2013.In: Purdue University Extension.

Mueller D 2016b. Corn disease loss estimates from the united states and ontario, canada - 2014.In: Purdue University Extension.

Mueller D 2017. Corn disease loss estimates from the united states and ontario, canada - 2016.In: Purdue University Extension.

Munkvold GP, White DG. 2016. Compendium of corn diseases: The American Phytopathological Society.

Nicholson RL, Hammerschmidt R. 1992. Phenolic-compounds and their role in disease resistance. Annual Review of Phytopathology 30: 369–389.

Olukolu BA, Wang GF, Vontimitta V, Venkata BP, Marla S, Ji J, Gachomo E, Chu K, Negeri A, Benson J, et al. 2014. A genome-wide association study of the maize hypersensitive defense response identifies genes that cluster in related pathways. PLoS Genet 10: e1004562.

Ono M, Takamura C, Sugita F, Masuoka C, Yoshimitsu H, Ikeda T, Nohara T. 2007. Two new steroid glycosides and a new sesquiterpenoid glycoside from the underground parts of trillium kamtschaticum. Chem Pharm Bull (Tokyo) 55: 551–556.

Panda P, Appalashetti M, Natarajan M, Chan-Park MB, Venkatraman SS, Judeh ZM. 2012a. Synthesis and antitumor activity of lapathoside d and its analogs. Eur J Med Chem 53: 1–12.

Panda P, Appalashetti M, Natarajan M, Mary CP, Venkatraman SS, Judeh ZM. 2012b.Synthesis and antiproliferative activity of helonioside a, 3’,4’,6’-tri-o-feruloylsucrose, lapathoside c and their analogs. Eur J Med Chem 58: 418–430.

Pasquet JC, Chaouch S, Macadre C, Balzergue S, Huguet S, Martin-Magniette ML, Bellvert F, Deguercy X, Thareau V, Heintz D, et al. 2014. Differential gene expression and metabolomic analyses of brachypodium distachyon infected by deoxynivalenol producing and non-producing strains of fusarium graminearum. Bmc Genomics 15.

Ponts N, Pinson-Gadais L, Boutigny AL, Barreau C, Richard-Forget F. 2011. Cinnamic-derived acids significantly affect fusarium graminearum growth and in vitro synthesis of type b trichothecenes. Phytopathology 101: 929–934.

Proctor RH, Hohn TM, Mccormick SP. 1995. Reduced virulence of gibberella zeae caused by disruption of a trichothecene toxin biosynthetic gene. Molecular Plant-Microbe Interactions 8: 593–601.

Schilmiller AL, Charbonneau AL, Last RL. 2012. Identification of a bahd acetyltransferase that produces protective acyl sugars in tomato trichomes. PNAS 109: 16377–16382.

Schmelz EA, Kaplan F, Huffaker A, Dafoe NJ, Vaughan MM, Ni XZ, Rocca JR, Alborn HT, Teal PE. 2011. Identity, regulation, and activity of inducible diterpenoid phytoalexins in maize. PNAS 108: 5455–5460.

Schnable PS, Ware D, Fulton RS, Stein JC, Wei F, Pasternak S, Liang C, Zhang J, Fulton L, Graves TA, et al. 2009. The b73 maize genome: Complexity, diversity, and dynamics. Science 326: 1112–1115.

Tautenhahn R, Bottcher C, Neumann S. 2008. Highly sensitive feature detection for high resolution lc/ms. Bmc Bioinformatics 9.

Vanden Wymelenberg AJ, Cullen D, Spear RN, Schoenike B, Andrews JH. 1997. Expression of green fluorescent protein in aureobasidium pullulans and quantification of the fungus on leaf surfaces. Biotechniques 23: 686–690.

Wang S, Basten, C.J., Zeng, Z-B. 2012. Windows qtl cartographer 2.5. Raleigh, NC: Dept. of Statistics, North Carolina State University.

Xiao J, Jin XH, Jia XP, Wang HY, Cao AZ, Zhao WP, Pei HY, Xue ZK, He LQ, Chen QG, et al. 2013. Transcriptome-based discovery of pathways and genes related to resistance against fusarium head blight in wheat landrace wangshuibai. Bmc Genomics 14.

Yan J, Aboshi T, Teraishi M, Strickler SR, Spindel JE, Tung CW, Takata R, Matsumoto F, Maesaka Y, McCouch SR, et al. 2015. The tyrosine aminomutase tam1 is required for beta-tyrosine biosynthesis in rice. Plant Cell 27: 1265–1278.

Yan L, Gao W, Zhang Y, Wang Y. 2008. A new phenylpropanoid glycosides from paris polyphylla var. Yunnanensis. Fitoterapia 79: 306–307.

Yang Q, He Y, Kabahuma M, Chaya T, Kelly A, Borrego E, Bian Y, El Kasmi F, Yang L, Teixeira P, et al. 2017. A gene encoding maize caffeoyl-coa o-methyltransferase confers quantitative resistance to multiple pathogens. Nat Genet 49: 1364–1372.

Ye JR, Guo YL, Zhang DF, Zhang N, Wang C, Xu ML. 2013. Cytological and molecular characterization of quantitative trait locus qrfg1, which confers resistance to gibberella stalk rot in maize. Molecular Plant-Microbe Interactions 26: 1417–1428.

Young TE, Meeley RB, Gallie DR. 2004. Acc synthase expression regulates leaf performance and drought tolerance in maize. Plant Journal 40: 813–825.

Zhang L, Liao CC, Huang HC, Shen YC, Yang LM, Kuo YH. 2008. Antioxidant phenylpropanoid glycosides from smilax bracteata. Phytochemistry 69: 1398–1404.

Zhang Y, He J, Jia LJ, Yuan TL, Zhang D, Guo Y, Wang YF, Tang WH. 2016. Cellular tracking and gene profiling of fusarium graminearum during maize stalk rot disease development elucidates its strategies in confronting phosphorus limitation in the host apoplast. Plos Pathogens 12.

Zhu JJ, Zhang CF, Zhang M, Wang ZT. 2006. Studies on chemical constituents in roots of rumex dentatus. Zhongguo Zhong Yao Za Zhi 31: 1691–1693.

